# Cross-species graph-embedding unmasks the ageing microenvironment as a key determinant of pancreatic cancer malignant cell biology and therapy response

**DOI:** 10.64898/2026.02.02.703350

**Authors:** Joaquín Araos Henríquez, Muntadher Jihad, Amir Jassim, Eloise G. Lloyd, Weike Luo, Judhell S. Manansala, Sneha Harish, Sara Pinto Teles, Priscilla S.W. Cheng, Gianluca Mucciolo, Wenlong Li, Marta Zaccaria, Debasmita Mukherjee, Rebecca Brais, Sally Mills, Paul M. Johnson, Mireia Vallespinos, Richard J. Gilbertson, Giulia Biffi

## Abstract

Pancreatic ductal adenocarcinoma (PDAC) has a dismal prognosis and is characterised by an extensive pro-tumorigenic stroma. Although most PDAC cases occur in older patients, the impact of ageing on malignant-stromal interactions and therapy response remains poorly understood. Here, we established orthotopically-grafted organoid-derived PDAC models across three murine age groups to characterise changes in the PDAC stroma and malignant cells with ageing. Cross-species analyses of tumour transcriptomes using a graph-embedding approach showed that integrating mouse models of different ages better captures the diversity of human PDAC, and that aged models more faithfully recapitulate the biology of older patients with PDAC. We also demonstrated that aged PDAC models have a more inflammatory stroma than that of younger tumours, shaping the malignant cell transcriptome. Finally, graph-embedding identified IRAK4 as a candidate therapeutic vulnerability in aged, but not young, KRAS- and p53-mutant PDAC, which we validated in preclinical drug studies. These findings highlight how ageing is a critical determinant of PDAC biology and associated therapeutic vulnerabilities, which should be an important consideration when designing disease models for preclinical development of precision therapies.

## Introduction

Pancreatic ductal adenocarcinoma (PDAC) is among the deadliest cancers, largely due to late diagnosis and therapy resistance^1^. PDAC includes an extensive and complex tumour microenvironment (TME) that is dominated by stromal cells that contribute to disease lethality by modulating therapy response and tumour progression^2,3^. Recent studies revealed that PDAC tumours contain variable amounts of heterogeneous populations of cancer-associated fibroblasts (CAFs) and other stromal cells, including macrophages and neutrophils^2,4,5^. Therefore, defining patterns of malignant cell-TME crosstalk in patients will be key for precision targeting.

Since most cases of PDAC are diagnosed after 70 years of age, the incidence of this cancer has increased dramatically over the last 30 years, concordant with increasing life expectancy^6^. Older patients with PDAC suffer from high levels of frailty and sarcopenia, and have worse post-resection survival rates^7^. Despite a strong association between old age and PDAC onset and clinical features, ageing remains largely overlooked in the design of preclinical models^17,18^. Indeed, PDAC is most frequently modelled in young mice (2-6 months old), which fail to recapitulate the aged physiological state present in most patients with PDAC^19^. Previous research with aged mouse models of various cancers has highlighted the impact of ageing on the TME, tumour growth, metastasis and therapy resistance^20–27^. Furthermore, comprehensive analysis of tissues across a broad range of ages has found that age-related transcriptional changes accumulate gradually and are missed in single-age studies^28^.

To address this gap, we established orthotopically-grafted organoid-derived PDAC models across three murine age groups (∼2-3, ∼12-13, and ∼18-20 months) to determine how an aged TME shapes PDAC malignant cell programs and therapeutically actionable vulnerabilities.

## RESULTS

### Ageing promotes a pro-inflammatory phenotype of the PDAC TME

Previous studies have shown that allografting malignant cells from a common source into mice of different ages can reveal age-dependent differences that are driven by the TME^21–25^; however, how differently aged pancreata impact the malignant cell phenotype remains unclear. Therefore, to determine how an aged TME impacts the biology of PDAC malignant cells, we transplanted PDAC organoids^29,30^ derived from ∼5-6-month-old KRAS- and p53-mutant KPC (*Kras^LSL-G12D/+^*; *Trp53^LSL-R172H/+^*; *Pdx1*-*Cre*) genetically engineered mouse models (GEMMs)^31^ into the pancreata of ∼2-3-month-old^32^ (hereon, moPDAC-young), ∼12-13-month-old (moPDAC-intermediate) and ∼18-20-month-old (moPDAC-old) C57BL/6J mice (**Fig. 1A**).

**Figure 1.**
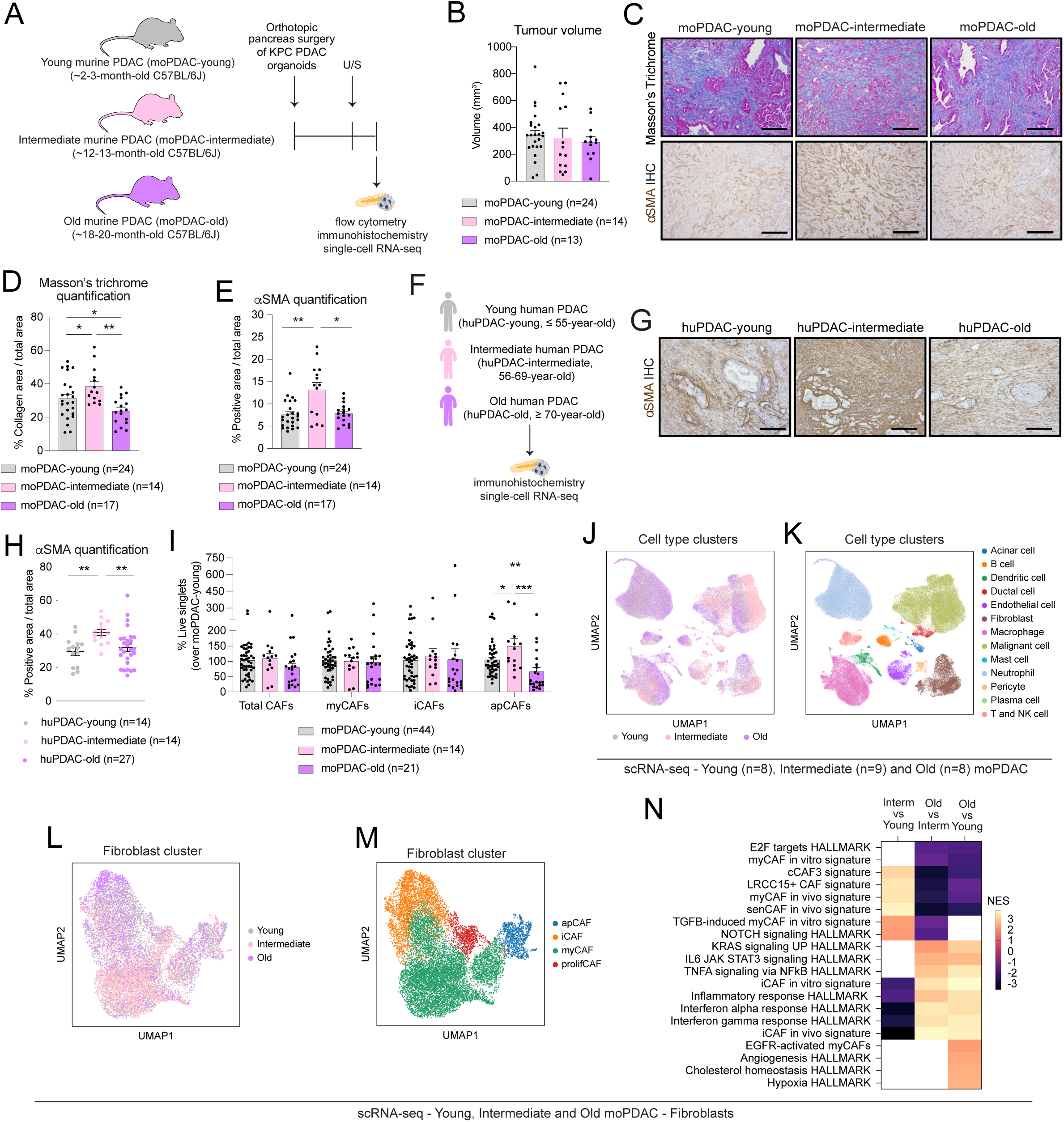
Ageing promotes an inflammatory CAF phenotype in PDAC mouse models. **(A)** Schematic of experimental design of analyses of orthotopically-grafted murine KPC (i.e., from *Kras^LSL-G12D/+^*, *Trp53^LSL-R172H/+^*, *Pdx1-Cre* GEMMs) organoid-derived pancreatic ductal adenocarcinoma (PDAC) models in young (∼2-3-month-old, moPDAC-young), intermediate (∼12-13-month-old, moPDAC-intermediate) and old (∼18-20-month-old, moPDAC-old) mice. U/S, ultrasound. **(B)** Volumes as measured by ultrasound-based imaging at the same time point of moPDAC-young, moPDAC-intermediate and moPDAC-old tumours. Results show mean ± SEM. No statistical difference was found, as calculated by Kruskal-Wallis test. **(C)** Representative Masson’s trichrome stains (top panels) and α-smooth muscle actin (αSMA, bottom panels) immunohistochemistry (IHC) stains in moPDAC-young, moPDAC-intermediate and moPDAC-old. Scale bars, 100 μm. **(D)** Quantification of Masson’s trichrome stain in moPDAC-young, moPDAC-intermediate and moPDAC-old. Results show mean ± SEM. *, *P adj* < 0.05; **, *P adj* < 0.01, Kruskal-Wallis test. **(E)** Quantification of αSMA stain in moPDAC-young, moPDAC-intermediate and moPDAC-old. Results show mean ± SEM. *, *P adj* < 0.05; **, *P adj* < 0.01, Kruskal-Wallis test. **(F)** Schematic of the young (≤ 55-year-old; hereon, huPDAC-young), intermediate (56–69-year-old; huPDAC-intermediate) and old (≥ 70-year-old; huPDAC-old) groups of human PDAC tumours analysed in this study. **(G)** Representative αSMA IHC stains in huPDAC-young, huPDAC-intermediate and huPDAC-old. Scale bars, 100 μm. **(H)** Quantification of αSMA stain in huPDAC-young, huPDAC-intermediate and huPDAC-old. Results show mean ± SEM. **, *P adj* < 0.01, Kruskal-Wallis test. **(I)** Flow cytometric analyses of total cancer-associated fibroblasts (CAFs; CD45^-^CD31^-^EpCAM^-^PDPN^+^), myofibroblastic CAFs (myCAFs; Ly6C^-^MHCII^-^), inflammatory CAFs (iCAF; Ly6C^+^MHCII^-^) and antigen-presenting CAFs (apCAFs; Ly6C^-^MHCII^+^) from live singlets in moPDAC-young, moPDAC-intermediate and moPDAC-old. Results show mean ± SEM. *, *P adj* < 0.05; **, *P adj* < 0.01; ***, *P adj* < 0.001, Kruskal-Wallis test. **(J-K)** Uniform Manifold Approximation and Projection (UMAP) plots of all cells from moPDAC-young (n=8), moPDAC-intermediate (n=9) and moPDAC-old (n=8) analysed by single-cell RNA-sequencing (scRNA-seq). Different age groups **(J)** or cell types **(K)** are colour coded. **(L-M)** UMAP plots of CAFs from moPDAC-young, moPDAC-intermediate and moPDAC-old analysed by scRNA-seq. Different ages **(L)** or CAF subsets **(M)** are colour coded. **(N)** Significantly upregulated and downregulated pathways identified by gene set enrichment analysis (GSEA) of CAFs from moPDAC-intermediate compared to moPDAC-young (left), moPDAC-old compared to moPDAC-intermediate (middle) and moPDAC-old compared to moPDAC-young (right), as assessed by MAST analysis from the scRNA-seq dataset. The *in vivo* iCAF and myCAF signatures were obtained from Elyada et al^34^. The *in vitro* iCAF and myCAF signatures were obtained from Öhlund et al^73^. The EGFR-activated CAF signature was obtained from Mucciolo and Araos Henríquez et al^32^. The myofibroblastic LRRC15^+^ CAF signature was obtained from Dominguez et al^101^, as published in Mucciolo and Araos Henríquez et al^32^. The myofibroblastic cCAF3 signature was obtained from McAndrews et al^102^, as published in Mucciolo and Araos Henríquez et al^32^. The senCAF *in vivo* signature was obtained from Belle et al^55^. Blank spaces indicate when gene signatures were not significantly enriched.

PDAC tumour growth rates were comparable across age groups (**Fig. 1B**). However, moPDAC-intermediate developed more liver metastases, whereas moPDAC-old exhibited more diaphragmatic metastases and greater weight loss than moPDAC-young (**Extended Data Fig. A-C**). Furthermore, levels of collagen and the myofibroblastic marker alpha smooth muscle actin (αSMA) varied with age in moPDAC, mirroring age-associated αSMA patterns observed in human (hu)PDAC-young (≤ 55-year-old^33^), huPDAC-intermediate (56-69-year-old) and huPDAC-old (≥ 70-year-old^33^) tumours (**Fig. 1C-H** and **Extended Data Fig. 1D-E**; **Supplementary Table 1**). Thus, the differently aged TME appeared to impact aspects of PDAC biology.

To further characterise age-related TME changes in PDAC, we leveraged a previously described flow cytometry strategy to quantify changes in CAF populations^32,34,35^. No significant difference in the total number of CAFs (PDPN^+^), myofibroblastic (my)CAFs (Ly6C^-^/MHCII^-^) or inflammatory (i)CAFs (Ly6C^+^/MHCII^-^) was seen among differently aged moPDACs (**Fig. 1I** and **Extended Data Fig. 1F-G**). However, antigen-presenting (ap)CAFs (Ly6C^-^/MHCII^+^) and the CD49E^+^ myCAF subset^35^ were significantly reduced in moPDAC-old relative to younger tumours (**Fig. 1I** and **Extended Data Fig. 1G**). In contrast to these relatively modest flow cytometry-defined changes in CAF composition, we observed marked changes in the single-cell RNA-sequencing (scRNA-seq) profiles of CAFs generated from moPDAC-young (n=8; n=45,208 cells), moPDAC-intermediate (n=9; n=41,456 cells) and moPDAC-old (n=8; n=47,914 cells) tumours (**Fig. 1J-N** and **Extended Data Fig. 1H-M**; **Supplementary Tables 2-4**). Pro-inflammatory gene sets (e.g., interferon [IFN], tumour necrosis factor [TNF] and JAK-STAT pathways), most of which have been previously associated with the iCAF phenotype^34,36–38^, were significantly enriched in scRNA-seq profiles of CAFs from moPDAC-old versus younger tumours (**Fig. 1N**). Furthermore, consistent with the increase in diaphragmatic metastases observed in moPDAC-old, the signature of an epithelial growth factor receptor (EGFR)-activated pro-metastatic myCAF subset^32^ was upregulated in moPDAC-old versus moPDAC-young (**Fig. 1N**). Thus, ageing appears to impact CAF phenotype in PDAC and promote a pro-inflammatory state.

Considering these changes, we next looked to see if PDAC immune cell populations varied with age. While the total numbers of T cells, B cells, Natural Killer (NK) cells and neutrophils did not vary significantly among differently aged moPDAC, moPDAC-old tumours contained significantly fewer macrophages overall but a higher proportion of major histocompatibility complex class II (MHCII)^+^ macrophages (**Fig. 2A-B** and **Extended Data Fig. 2A-D**). Furthermore, analysis of scRNA-seq profiles using previously defined markers of tumour-associated macrophages (TAMs)^4,35^ identified transcriptomic changes across age groups, including an increase in interferon-primed TAM (IFN-TAM) markers and associated pathways in moPDAC-old compared to younger tumours (**Fig. 2C-F** and **Extended Data Fig. 2E**; **Supplementary Tables 2-4**). Gene signatures of neutrophil states recently described in PDAC^5,35^ were also altered with age, with the tumour-promoting T3 neutrophil signature being significantly upregulated in moPDAC-old compared to both younger tumour groups and the T2 mature neutrophil signature being significantly upregulated in moPDAC-old compared to moPDAC-young (**Fig. 2G-I** and **Extended Data Fig. 2F-K**; **Supplementary Tables 2-4**).

**Figure 2.**
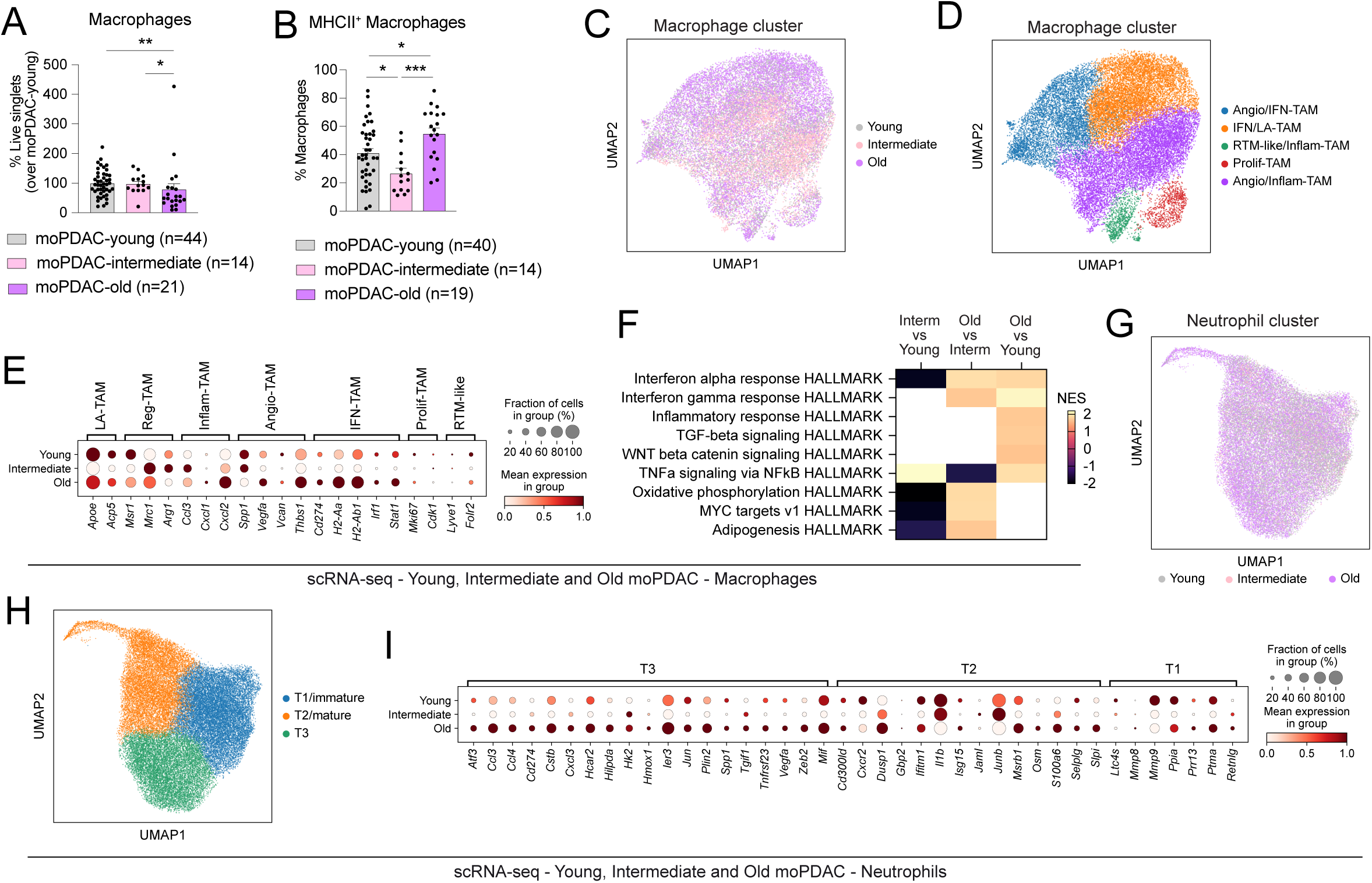
Ageing shapes immune cell composition in PDAC mouse models. **(A)** Flow cytometric analysis of macrophages (CD45^+^CD11b^+^Gr1^-^F4/80^+^) from live singlets in moPDAC-young, moPDAC-intermediate and moPDAC-old. Results show mean ± SEM. *, *P adj* < 0.05; **, *P adj* < 0.01, Kruskal-Wallis test. **(B)** Flow cytometric analysis of major histocompatibility complex class II (MHCII)^+^ macrophages from the parental gate in moPDAC-young, moPDAC-intermediate and moPDAC-old. Results show mean ± SEM. *, *P adj* < 0.05; ***, *P adj* < 0.001, Kruskal-Wallis test. **(C-D)** UMAP plots of macrophages from moPDAC-young (n=8), moPDAC-intermediate (n=9) and moPDAC-old (n=8) analysed by scRNA-seq. Different ages **(C)** or subsets **(D)** are colour coded. **(E)** Dot plot visualization of the scaled average expression of macrophage subset markers in macrophages from moPDAC-young, moPDAC-intermediate and moPDAC-old analysed by scRNA-seq. The colour intensity represents the expression level, and the size of the dots represents the percentage of expressing cells. **(F)** Selected significantly upregulated and downregulated pathways identified by GSEA of macrophages from moPDAC-intermediate compared to moPDAC-young (left), moPDAC-old compared to moPDAC-intermediate (middle) and moPDAC-old compared to moPDAC-young (right), as assessed by MAST analysis from the scRNA-seq dataset. Blank spaces indicate when gene signatures were not significantly enriched. **(G-H)** UMAP plots of neutrophils from moPDAC-young, moPDAC-intermediate and moPDAC-old analysed by scRNA-seq. Different ages **(G)** or subsets **(H)** are colour coded. **(I)** Dot plot visualization of the scaled average expression of neutrophil subset markers in neutrophils from moPDAC-young, moPDAC-intermediate and moPDAC-old analysed by scRNA-seq. The colour intensity represents the expression level, and the size of the dots represents the percentage of expressing cells.

Together, these findings reveal significant and complex age-related changes in the TME of aged mouse models of PDAC, most evident within tumour transcriptomes, that include the reprogramming of fibroblasts and immune cells towards a pro-inflammatory state.

### Graph-embedding reveals age-dependent acquisition of inflammatory functions in PDAC

We recently reported that measuring changes in the co-expression context of genes across single cell transcriptomes of different tumour states – using an approach we termed *RE*sistance through *CO*ntext *DR*ift (RECODR)^39^, can unmask biological changes not detected by conventional analyses (e.g., differential gene expression) or other graph network-based approaches (e.g., Weighted Gene Co-expression Network Analysis, WGCNA)^39^. Briefly, RECODR generates gene co-expression graph networks (GNs) from scRNA-seq profiles, organising genes with correlated expression, and therefore potentially related function, into distinct communities. RECODR then compares GNs across conditions (e.g., old vs young PDAC) to measure the degree to which each gene has changed its co-expression context between these different tissue states. RECODR assigns each gene a ‘context drift score’ (CDS), quantifying how much its co-expression neighbourhood changes between the tissue states. Genes with the highest CDS (i.e., that have shifted their transcriptome context the most) are predicted to contribute the most to the change in biological state^39^. Importantly, extensive benchmarking showed that this approach detected biologically and therapeutically meaningful information that was invisible to existing analytical approaches and was independent of differences in gene expression levels^39^.

GNs generated by RECODR from scRNA-seq profiles of moPDAC-young (hereon, GN^moPDAC-young^; n=8,952 genes), moPDAC-intermediate (GN^moPDAC-interm^; n=6,660 genes) and moPDAC-old (GN^moPDAC-old^; n=8,940 genes) were similar (**Fig. 3A**; **Supplementary Table 5**). Pathway enrichment analysis revealed that each GN contained four major community types that were enriched for genes associated with similar functions: cell signalling/interactions, metabolism, extracellular matrix (ECM) and inflammatory (**Fig. 3A-B**; **Supplementary Table 5**). Around one third of genes on the GN^moPDAC-old^ (n=3,233/8,940) were located in the same community type on both the GN^moPDAC-young^ and GN^moPDAC-interm^ (we termed these ‘totally retained’ genes), and another one third of GN^moPDAC-old^ genes (n=2,681/8,940) were located in the same community type on either the GN^moPDAC-young^ or the GN^moPDAC-interm^ (we termed these ‘partially retained’ genes; **Fig. 3B**; **Supplementary Table 5**). These findings indicate that substantial aspects of tumour biology are shared among differently aged moPDAC models.

**Figure 3.**
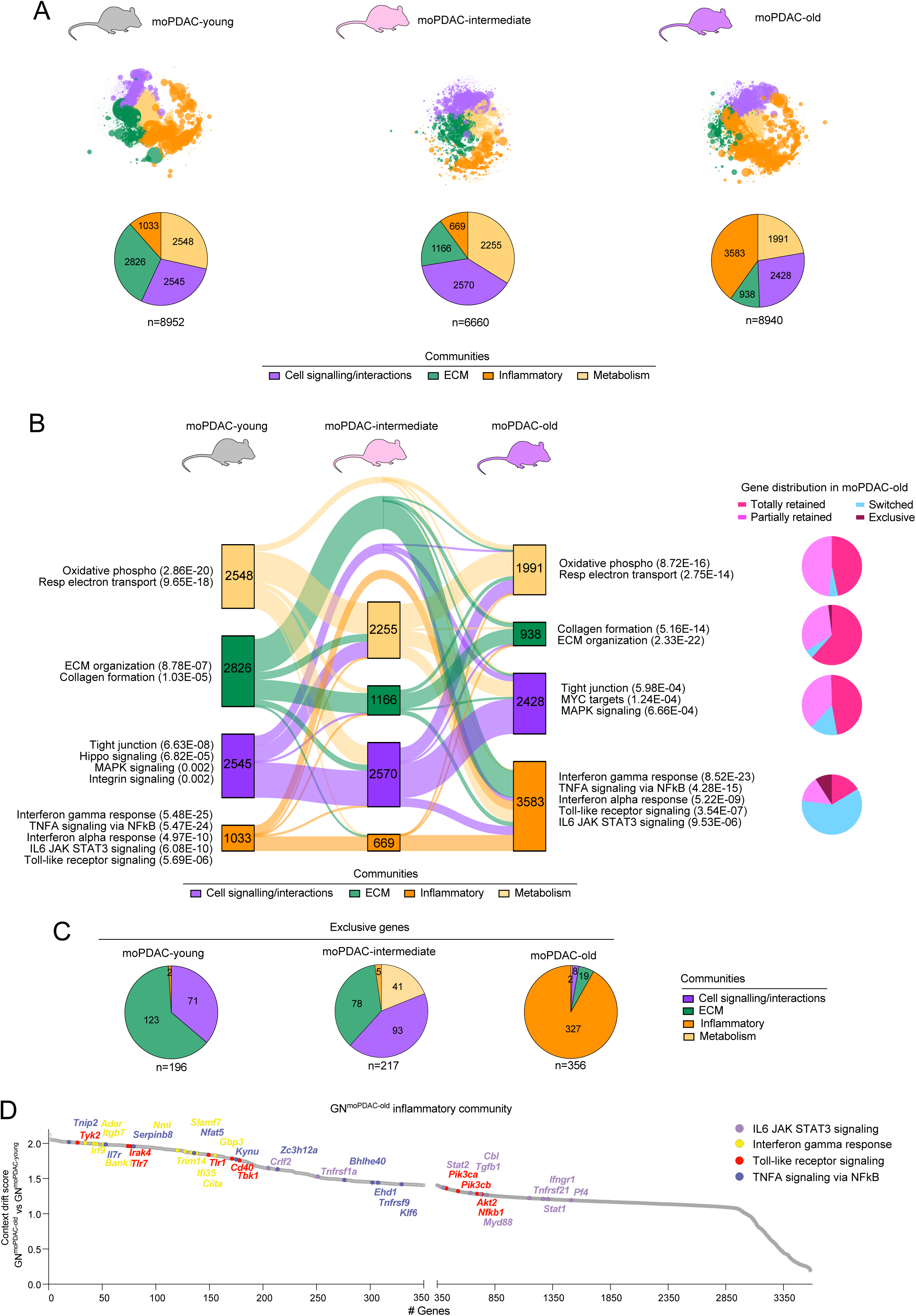
Graph-embedding reveals age-dependent acquisition of inflammatory functions in PDAC mouse models. **(A)** *RE*sistance through *CO*ntext *DR*ift (RECODR) graph networks (GNs; top) and pie charts (bottom) of gene communities generated from scRNA-seq datasets of moPDAC-young (n=8; hereon, GN^moPDAC-young^; left; n=8,952 genes; n=45,208 cells), moPDAC-intermediate (n=9; GN^moPDAC-interm^; middle; 6,660 genes; n=41,456 cells) and moPDAC-old (n=8; GN^moPDAC-old^; right; n=8,940 genes; n=47,914 cells). **(B)** Left: alluvial plot of gene locations in communities from GN^moPDAC-young^, GN^moPDAC-interm^ and GN^moPDAC-old^. Enriched pathways in relevant gene communities (i.e., with > 35 genes) are shown with p adj < 0.05 in brackets. Each box represents a community, and the size of each box is normalised to the number of genes (indicated in the boxes) per community. The size of each connecting ribbon between two boxes is proportional to the number of genes that are found in that pair of communities. Right: pie charts report the proportion of ‘exclusive, ‘switched’, ‘partially retained’ and ‘totally retained’ genes in the corresponding communities of GN^moPDAC-old^. **(C)** Pie charts showing the distribution within communities of genes ‘exclusive’ to GN^moPDAC-young^ (left, n=196), GN^moPDAC-interm^ (middle, n=217) and GN^moPDAC-old^ (right, n=356). **(D)** Ranked context drift score (CDS) of inflammatory genes on the GN^moPDAC-old^ compared to GN^moPDAC-young^. Highlighted genes are the top 10 in terms of highest CDS for the four indicated colour-coded pathways based on pathway enrichment analysis of inflammatory genes on GN^moPDAC-old^.

However, RECODR also detected marked differences in gene co-expression patterns among the three different GNs. Of note, ‘totally retained’ genes in the GN^moPDAC-old^ were unevenly distributed among the four communities. Fewer genes were ‘totally retained’ in the inflammatory community (16%, n=583/3,583; e.g., antigen presentation regulators *H2-Ab1*, *H2-Aa*, *Cd74*) compared to the ECM (62%, n=577/938; e.g., transforming growth factor beta [TGF-β] signalling components *Tgfb2*, *Tgfb3*), metabolism (47%, n=930/1,991; e.g., mevalonate pathway components *Pmvk*, *Hmgcr*) and cell signalling/interactions (47%, n=1,143/2,428; e.g., integrins *Itga6*, *Itgb4*) communities (chi-square test, *P* < 0.001; **Fig. 3B**, **Extended Data Fig. 3A**; **Supplementary Table 5**). Furthermore, 30% of GN^moPDAC-old^ genes (n=2,670/8,940) also present on the GN^moPDAC-young^ and/or GN^moPDAC-interm^ were located in different community types compared to both younger ages (we termed these ‘switched’ genes; **Fig. 3B**; **Supplementary Table 5**). These genes included mitochondria-related genes (*Bnip1*, *Bnip3l*, *Higd1a*) in the GN^moPDAC-young^ ECM community that ‘switched’ to the GN^moPDAC-old^ inflammatory community. Quantitatively, significantly more GN^moPDAC-old^ genes were ‘switched’ in the inflammatory community (61%, n=2,170/3,583) compared to the cell signalling/interactions (16%, n=356/2,428), ECM (5%, n=44/938) and metabolism (5%, n=100/1,991) communities (chi-square test, *P* < 0.001; **Fig. 3B**; **Supplementary Table 5**). Finally, among genes that were restricted exclusively to a single GN (we termed these ‘exclusive’ genes), those on the GN^moPDAC-old^ were significantly enriched within the inflammatory community. Indeed, 92% (n=327/356) of GN^moPDAC-old^ ‘exclusive’ genes were in the inflammatory community, compared with only 2% (n=5/217) and 1% (n=2/196) of ‘exclusive’ genes in this community type on the GN^moPDAC-interm^ and GN^moPDAC-young^, respectively (chi-square test, *P* < 0.001; **Fig. 3B-C** and **Extended Data Fig. 3B**). GN^moPDAC-old^ ‘exclusive’ genes further enriched the GN^moPDAC-old^ inflammatory community for regulators of toll-like receptor (TLR; *Tlr1, Tlr7, Irak4*), TNF (*Il7r*, *Myd88*, *Tnfrsf26*), IFN (*Irf9*, *Ifi35*, *Ciita*) and JAK-STAT (*Stat5b*, *Tyk2*) signalling, with pro-inflammatory genes from these pathways displaying high CDS values (**Fig. 3D**). Conversely, ECM ‘exclusive’ genes dropped with age, with only 5% (n=19/356) in the GN^moPDAC-old^ ECM community compared to 36% (n=78/217) and 63% (n=123/196) of genes in this community on the GN^moPDAC-interm^ and GN^moPDAC-young^, respectively (chi-square test, *P* < 0.001; **Fig. 3C**). Thus, in keeping with our conventional scRNA-seq analysis, RECODR also pinpointed age-related remodelling of PDAC transcriptomes towards a pro-inflammatory state.

Mapping inflammatory community genes back to individual scRNA-seq profiles revealed enriched expression in macrophages, neutrophils and dendritic cells (**Fig. 1K** and **Extended Data Fig. 3C**). Notably, in agreement with prior studies of brain and breast cancers^39^, RECODR CDS resolved transcriptomic differences between GN^moPDAC-old^ and GN^moPDAC-young^ more effectively than differential expression analysis, and highlighted generally higher CDS values among ‘exclusive’ genes (**Extended Data Fig. 3D**; **Supplementary Tables 2** and **5**). Focusing on the inflammatory community, CDS clearly resolved genes within this community, and scored TLR, TNF, IFN and JAK-STAT signalling mediators particularly highly (**Extended Data Fig. 3E**; **Supplementary Table 5**). Indeed, while 71% (n=2,552/3,583) of genes in the GN^moPDAC-old^ inflammatory community were absent from the GN^moPDAC-young^ inflammatory community, only 125 genes were significantly differentially expressed between immune cells (comprising macrophages, neutrophils and dendritic cells) in moPDAC-old versus moPDAC-young (average Log2-fold change [avg Log2FC] > 0.05; false discovery rate [FDR] <0.05; **Extended Data Fig. 3F**; **Supplementary Table 5**).

Thus, RECODR pinpointed profound age-related transcriptomic remodelling in moPDAC tumours, marked by acquisition of inflammatory functions, comprising the TLR, TNF, IFN and JAK-STAT pathways, suggesting these might contribute the most to the increased pro-inflammatory phenotype of old PDAC.

### Old PDAC mouse models capture age-specific inflammatory changes of human PDAC

To determine whether the differences observed among moPDAC-young, -intermediate and -old models reflect age-dependent changes in human PDAC, we analysed previously reported^40^ scRNA-seq profiles of young (≤ 55-year-old; n=11; n=21,769 cells), intermediate (56-69-year-old; n=26; n=41,292 cells) and old (≥ 70-year-old^33^; n=18; n=41,677 cells) huPDAC tumours (**Supplementary Table 6**). Conventional analyses of huPDAC scRNA-seq profiles identified similar cell types to those observed in moPDAC (**Extended Data Fig. 4A-E**). Furthermore, RECODR-generated GNs of young (hereon, GN^huPDAC-young^; n=3,980 genes), intermediate (GN^huPDAC-interm^; n=7,006 genes) and old (GN^huPDAC-old^; n=8,460 genes) huPDAC contained gene communities broadly similar to those of our organoid-derived moPDAC models in both numbers and functions, including cell signalling/interactions, metabolism, ECM and inflammatory (**Fig. 4A-B** and **Extended Data 4F**; **Supplementary Table 7**).

**Figure 4.**
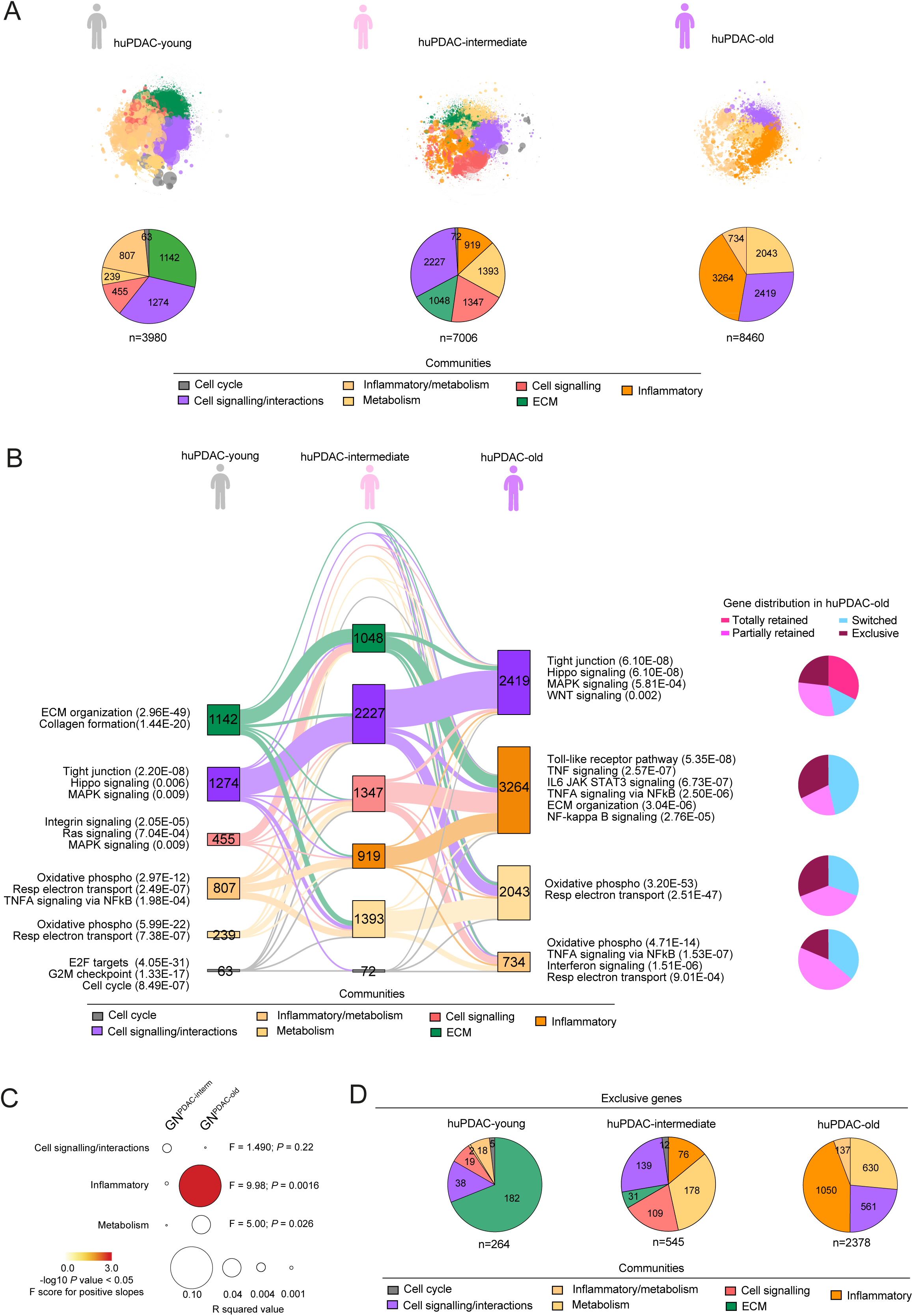
Old PDAC mouse models capture age-specific changes of human PDAC. **(A)** GNs (top) and pie charts (bottom) of gene communities generated from scRNA-seq datasets of huPDAC-young (n=11; hereon, GN^huPDAC-young^; left; n=3,980 genes; n=21,769 cells), huPDAC-intermediate (n=26; GN^huPDAC-interm^; middle; n=7,006 genes; n=41,292 cells) and huPDAC-old (n=18; GN^huPDAC-old^; right; n=8,460 genes; n=41,677 cells). scRNA-seq data are from Chijimatsu et al^40^. **(B)** Left: alluvial plot of gene locations in communities from GN^huPDAC-young^, GN^huPDAC-interm^ and GN^huPDAC-old^. Enriched pathways in relevant gene communities (i.e., with > 35 genes) are shown with p adj < 0.05 in brackets. Each box represents a community, and the size of each box is normalised to the number of genes (indicated in the boxes) per community. The size of each connecting ribbon between two boxes is proportional to the number of genes that are found in that pair of communities. Right: pie charts report the proportion of ‘exclusive, ‘switched’, ‘partially retained’ and ‘totally retained’ genes in the corresponding communities of GN^huPDAC-old^. **(C)** Dot plot showing cross-species analysis of linear regression between CDS of specific gene communities within GN^moPDAC^ and GN^huPDAC^. Cross-species comparisons across common genes between GN^PDAC-old^ and GN^PDAC-young^ (GN^PDAC-old^) and across common genes between GN^PDAC-interm^ and GN^PDAC-young^ (GN^PDAC-interm^) are shown. The colour intensity represents the statistical significance of the F-test results (for positive slops and *P* < 0.05), and the size of the dots represents the R squared value. F-test results are shown as comparisons between each pair of slopes. **(D)** Pie charts showing the distribution within communities of genes ‘exclusive’ to GN^huPDAC-young^ (left, n=264), GN^huPDAC-interm^ (middle, n=545) and GN^huPDAC-old^ (right, n=2,378).

As in moPDAC, RECODR identified age-related differences in gene co-expression patterns among huPDAC tumours that were not detected by differential expression analysis (**Extended Data Fig. 4G**; **Supplementary Table 7**). Furthermore, cross-species analysis of gene context drift between young and intermediate, and young and old PDACs, revealed that moPDAC-old more closely mirrored the gene context drift of huPDAC-old than moPDAC-intermediate mirrored the gene context drift of huPDAC-intermediate (**Extended Data Fig. 4H**). Of note, this cross-species correlation in CDS was strongest for genes in the inflammatory communities of GN^huPDAC-old^ and GN^moPDAC-old^ (**Fig. 4C**).

Changes in the size and gene enrichment of the GN^huPDAC-old^ inflammatory community relative to younger human tumours were similar to those observed in moPDAC: (i) 44% of GN^huPDAC-old^ ‘exclusive’ genes (n=1,050/2,378) were in the inflammatory community compared to only 14% (n=76/545) and 0% (n=0/264) of ‘exclusive’ genes in this community type on the GN^huPDAC-interm^ and GN^huPDAC-young^, respectively (chi-square test, *P* < 0.001; **Fig. 4D**); and (ii) GN^huPDAC-old^ inflammatory ‘exclusive’ genes were significantly enriched for TLR (*IRAK4*, *MAP3K7*) and TNF (*MAPK1*, *CREB1*) signalling (**Supplementary Table 7**). Finally, inflammatory ‘exclusive’ genes on the GN^moPDAC-old^, but not on GN^moPDAC-interm^ or GN^moPDAC-young^, significantly overlapped with GN^huPDAC-old^ inflammatory ‘exclusive’ genes (**Extended Data Fig. 4I**, *P* < 0.001, hypergeometric test; **Supplementary Table 8**). To evaluate whether moPDAC-old more faithfully captured the age-specific biology of huPDAC-old also compared to widely used KPC GEMMs (< 1-year-old), we leveraged RECODR to generate a GN from previously published scRNA-seq profiles (n=4; n=15,075 cells) of KPC GEMM-derived PDACs^34^ (hereon, GN^moPDAC-GEMM^; n=5,137 genes; **Extended Data Fig. 4J-K**; **Supplementary Table 9**). Of note, also in this analysis, inflammatory genes ‘exclusive’ to GN^moPDAC-old^ compared to GN^moPDAC-GEMM^ significantly overlapped with inflammatory genes ‘exclusive’ to GN^huPDAC-old^ (**Extended Data Fig. 4L-M**, *P* < 0.001, hypergeometric test; **Supplementary Table 9**). Additionally, genes in other communities (e.g., metabolism and cell signalling/interactions) and ‘exclusive’ to GN^moPDAC-old^ compared to GN^moPDAC-GEMM^ more significantly overlapped with GN^huPDAC-old^ and GN^huPDAC-interm^ ‘exclusive’ genes in the related community type (**Extended Data Fig. 4M**; **Supplementary Table 9**).

Together, these findings indicate that aged mouse models better recapitulate age-dependent gene expression patterns observed in the majority of PDAC patients (i.e., those ≥ 70-year-old). These data further highlight increased inflammatory functions in both murine and human old PDACs relative to younger tumours, suggesting these to be potential age-specific therapeutic vulnerabilities.

### Cross-species graph-embedding highlights the ageing TME as a key determinant of PDAC malignant cell biology

Having charted transcriptome evolution across all cells within PDAC tumours, we assessed how an aged TME might influence malignant cells. To do this, we first identified murine epithelial malignant cell profiles using InferCNV and known markers (**Extended Data Fig. 5A** and **Extended Data Fig. 1K-L**). These accounted for 26% (n=11,603/45,208), 47% (n=19,522/41,456) and 17% (n=8,358/47,914) of scRNA-seq profiles in moPDAC-young, - intermediate and -old, respectively. In keeping with our analyses of all PDAC cell types, gene set enrichment analysis (GSEA) revealed significant upregulation of pro-inflammatory pathways, including IFN, TNF and JAK/STAT signalling in moPDAC-old compared to moPDAC-young malignant cells (**Extended Data Fig. 5B; Supplementary Table 10**). We then looked to see if moPDAC malignant cells included subpopulations that might drive these transcriptional changes. Three malignant cell subsets were detected across all age groups, and were enriched in mesenchymal (subset 1), inflammatory (subset 2) or proliferative (subset 3) markers (**Fig. 5A** and **Extended Data Fig. 5C-D**). We found that the relative proportion of these malignant cell subsets did not vary with ageing, while inflammatory pathway genes were enriched to varying degrees in moPDAC-old malignant cell subsets compared to moPDAC-young (**Fig. 5B-C**; **Supplementary Table 10**).

**Figure 5.**
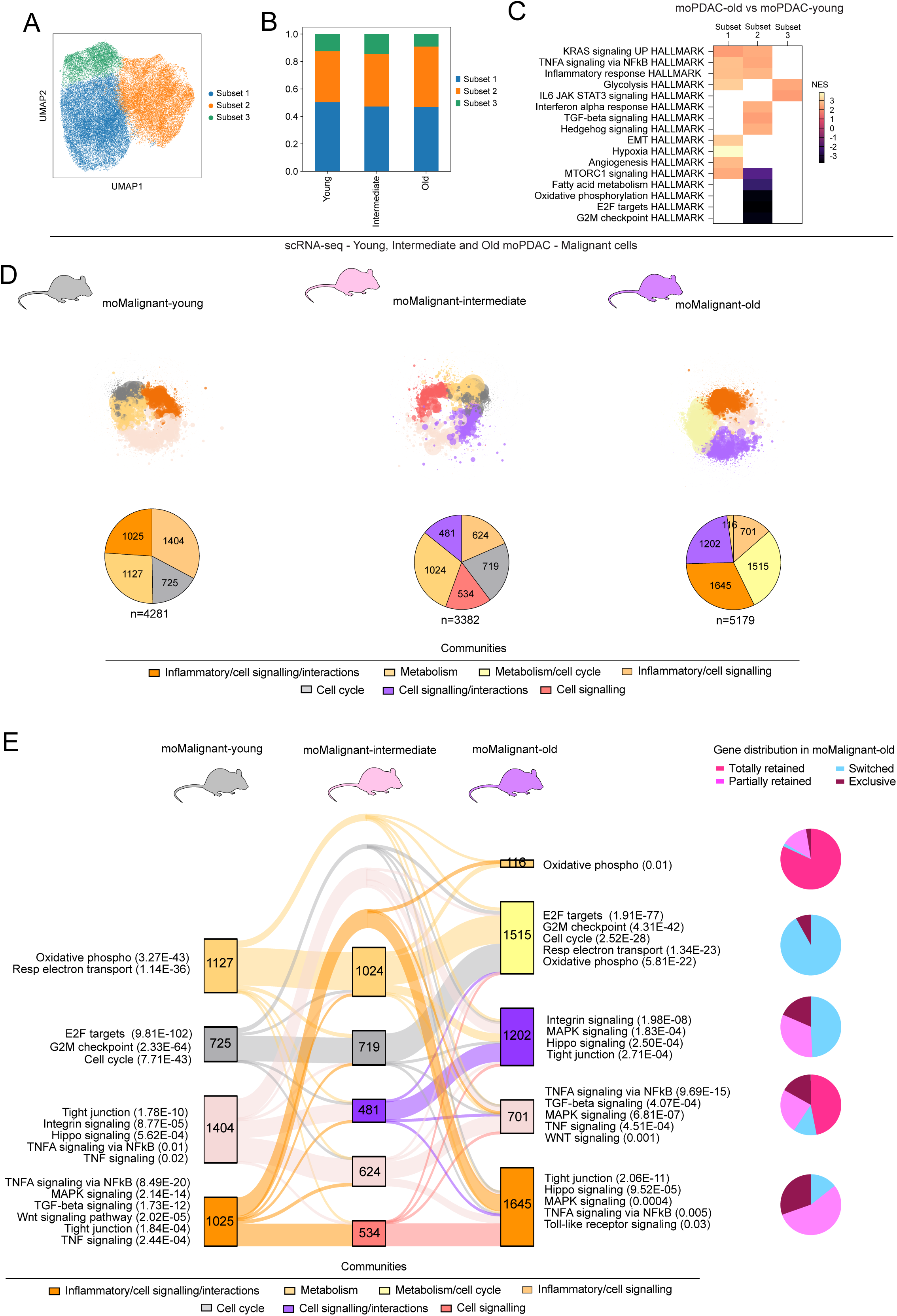
The aged TME promotes acquisition of pro-inflammatory functions in murine PDAC malignant cells. **(A)** UMAP plot of malignant cells from moPDAC-young, moPDAC-intermediate and moPDAC-old analysed by scRNA-seq. Different subsets are colour coded. **(B)** Subset contribution in malignant cells from moPDAC-young, moPDAC-intermediate and moPDAC-old represented as bar plot showing proportions of the different subsets in each condition. **(C)** Significantly upregulated and downregulated pathways identified by GSEA of each malignant cell subpopulation from moPDAC-old compared to moPDAC-young, as assessed by MAST analysis from the scRNA-seq dataset. Blank spaces indicate when gene signatures were not significantly enriched. **(D)** GNs (top) and pie charts (bottom) of gene communities in malignant cells from moPDAC-young (hereon, GN^moMalign-young^; left; n=4,281 genes; n=11,603 cells), moPDAC-intermediate (GN^moMalign-interm^; middle; n=3,382 genes; n=19,522 cells) and moPDAC-old (GN^moMalign-old^; right; n=5,179 genes; n=8,358 cells). **(E)** Left: alluvial plot of gene locations in communities from GN^moMalign-young^, GN^moMalign-interm^ and GN^moMalign-old^. Enriched pathways in relevant gene communities (i.e., with > 35 genes) are shown with p adj < 0.05 in brackets. Each box represents a community, and the size of each box is normalised to the number of genes (indicated in the boxes) per community. The size of each connecting ribbon between two boxes is proportional to the number of genes that are found in that pair of communities. Right: pie charts report the proportion of ‘exclusive’, ‘switched’, ‘partially retained’ and ‘totally retained’ genes in the corresponding communities of GN^moMalign-old^.

To further understand differences among moPDAC malignant cell transcriptomes that may be driven by differently aged TME, we applied RECODR to generate GNs of young (hereon, GN^moMalign-young^; n=4,281 genes), intermediate (GN^moMalign-interm^; n=3,382 genes) and old (GN^moMalign-old^; n=5,179 genes) moPDAC malignant cells (**Fig. 5D-E**; **Supplementary Table 11**). RECODR identified communities related to metabolism, inflammation, cell signalling and cell interactions (**Fig. 5E**). Notably, as with GNs of all moPDAC cells, the number of ‘exclusive’ genes present on malignant cell-only GNs increased with age (n=120 in GN^moMalign-young^; n=150 in GN^moMalign-interm^; n=970 in GN^moMalign-old^), and were markedly located in the GN^moMalign-old^ inflammatory/cell signalling/interactions community (52%, n=500/970; **Fig. 5E** and **Extended Data Fig. 5E-F**). Finally, RECODR CDS again resolved gene expression patterns between moPDAC-old and moPDAC-young malignant cells better than differential gene expression analysis, and highlighted ‘exclusive’ genes having among the highest CDS values (**Extended Data Fig. 5G**; **Supplementary Table 11**). Thus, ageing promotes a pro-inflammatory phenotype in moPDAC-old also in malignant cells. Moreover, since a common source of PDAC malignant cells was grafted into our mouse models, the age of the host TME appears to directly impact the transcriptome of malignant cells.

We next evaluated whether these changes in moPDAC malignant cell transcriptomes recapitulated those in huPDAC malignant cells. Using inferCNV and epithelial cell marker expression, we identified the malignant cells within scRNA-seq profiles of huPDAC-young (n=3,479/21,769), -intermediate (n=11,295/41,292) and -old (n=11,860/41,677; **Extended Data Fig. 4C-D**). Like for moPDAC malignant cells, RECODR identified communities related to metabolism, inflammation, cell signalling and cell interactions in GNs from young (hereon, GN^huMalign-young^; n=3,310 genes), intermediate (GN^huMalign-interm^; n=6,681 genes) and old (GN^huMalign-old^; n=9,172 genes) huPDAC malignant cells (**Fig. 6A-B**; **Supplementary Table 12**). Additionally, as observed in the GNs of moPDAC malignant cells, ‘exclusive’ genes in huPDAC malignant cell GNs increased with age (n=191 in GN^huMalign-young^; n=915 GN^huMalign-interm^; n=3,212 GN^huMalign-old^) and were enriched in inflammatory communities (**Extended Data Fig. 6A-B**). Notably, genes that were ‘exclusive’ to the GN^moMalign-old^ and GN^huMalign-old^ inflammatory/cell signalling-related communities overlapped significantly (**Fig. 6C**; *P* < 0.001, hypergeometric test; **Supplementary Table 13**). Consistent with our comparisons of ‘exclusive’ genes between GN^moPDAC-old^ and GN^huPDAC-old^, this degree of overlap was not seen with younger mouse models (**Fig. 6C** and **Extended Data Fig. 4I**).

**Figure 6.**
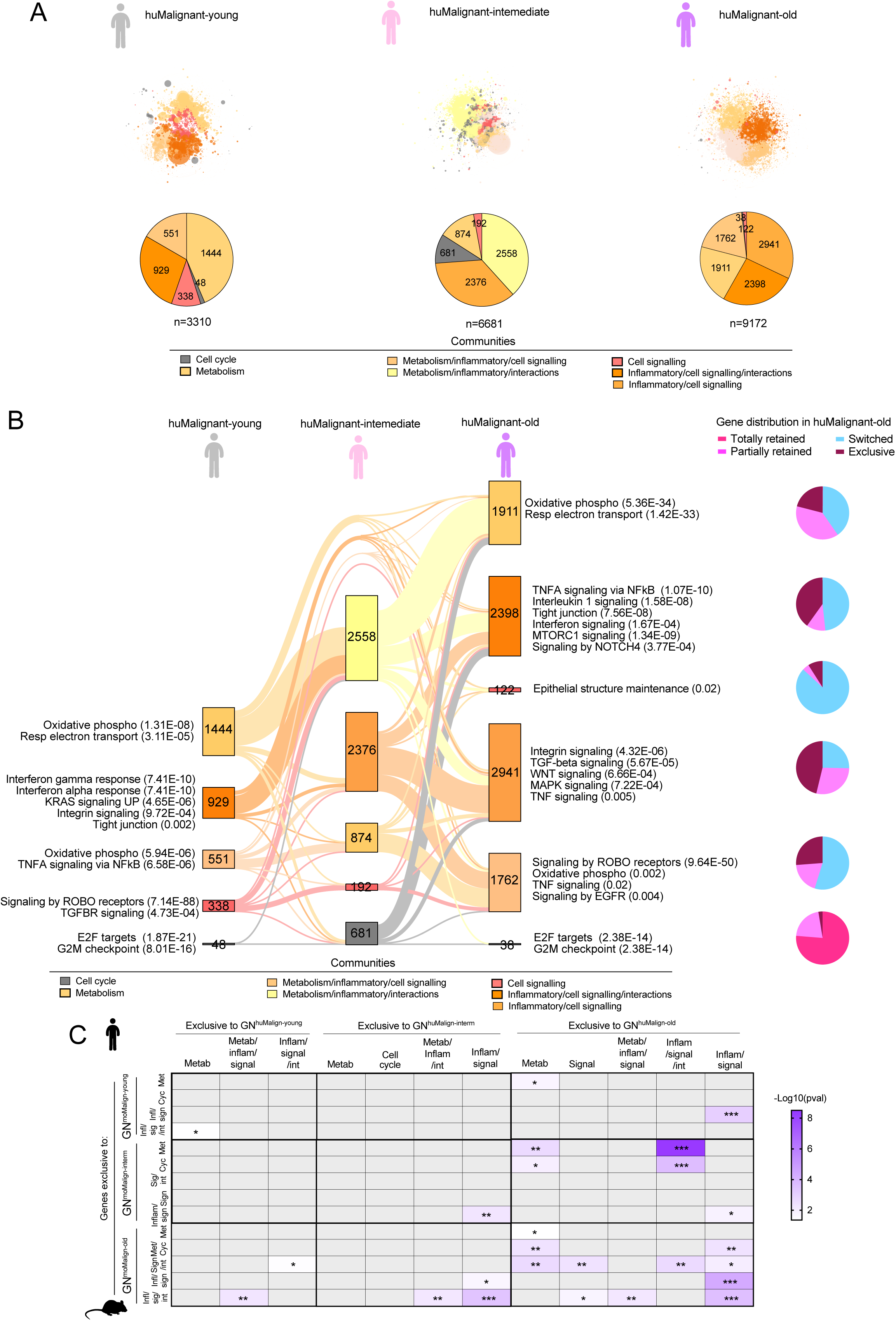
Cross-species graph-embedding highlights the aged TME as a key determinant of PDAC malignant cell biology. **(A)** GNs (top) and pie charts (bottom) of gene communities in malignant cells from huPDAC-young (hereon, GN^huMalign-young^; left; n=3,310 genes; n=3,479 cells), huPDAC-intermediate (GN^huMalign-interm^; middle; n=6,681 genes; n=11,295 cells) and huPDAC-old (GN^huMalign-old^; right; n=9,172 genes; n=11,860 cells). **(B)** Left: alluvial plot of gene locations in communities from malignant cell GNs of young, intermediate, and old human PDAC. Enriched pathways in relevant gene communities (i.e., with > 35 genes) are shown with p adj < 0.05 in brackets. Each box represents a community, and the size of each box is normalised to the number of genes (indicated in the boxes) per community. The size of each connecting ribbon between two boxes is proportional to the number of genes that are found in that pair of communities. Right: pie charts report the proportion of ‘exclusive’, ‘switched’, ‘partially retained’ and ‘totally retained’ genes in the corresponding communities of GN^huMalign-old^. **(C)** Heatmap showing the overlap of ‘exclusive’ genes in each community on GN^moMalign-young^, GN^moMalign-interm^ and GN^moMalign-old^ with ‘exclusive’ genes in each community on GN^huMalign-young^, GN^huMalign-interm^ and GN^huMalign-old^. Grey spaces indicate not-significant overlaps. Significance was defined by hypergeometric test with representation factor > 1; * *P* < 0.05, ** *P* < 0.01, *** *P* < 0.001; denominator = 17,094 orthologs.

Together, these data show that old PDAC mouse models more faithfully capture age-specific pro-inflammatory features of huPDAC malignant cells. Since our moPDAC models originated from a common source of organoids, these findings suggest that an aged TME directly remodels PDAC malignant cell transcriptomes, shaping biology and therapeutic vulnerabilities.

### IRAK4 activity is a therapeutic vulnerability of old PDAC

Tumours expressing similar levels of a drug target can respond differently to its therapeutic inhibitor, making context prediction a major challenge in cancer therapy^41^. To address this, we again leveraged RECODR, which was originally developed to predict the gene context in which a therapy is likely to succeed or fail^39^. We hypothesised that RECODR might predict therapeutic vulnerabilities specific to PDAC tumours of different ages. As a first step to test this hypothesis, we sought to identify genes predicted by RECODR as context-specific therapeutic targets in PDAC. Since inflammation-related communities were most prominent in moPDAC-old and huPDAC-old relative to younger tumours, and showed significant cross-species overlap in both all cell- and malignant cell-derived GNs (**Extended Data Fig. 4I** and **Fig. 6C**), we focused on genes within these communities. Furthermore, because genes with high CDS are predicted by RECODR to contribute substantially to changes in biological state^39^, and genes ‘exclusive’ to a GN exhibited among the highest CDS values (**Extended Data Fig. 3D** and **5G**), we focused on genes that were ‘exclusive’ to moPDAC-old and huPDAC-old GNs when compared with their respective young GNs. We prioritised candidate targets by selecting genes encoding proteins with favourable druggability features, including: (i) high drug-gene specificity based on Drug-Gene interaction database (DGIdb) scores > 1^42^; (ii) high predicted ligand druggability based on canSAR.ai protein annotation tool (CPAT) ligand druggability scores (LDS) > 1^43^; and (iii) the availability of selective inhibitors based on CPAT annotations^43^. This strategy identified two druggable orthologs shared between moPDAC-old and huPDAC-old inflammatory communities, and six druggable orthologs shared between inflammatory-related communities in moPDAC-old and huPDAC-old malignant cells (**Fig. 7A**; **Supplementary Table 14**). Among these candidates, we selected *IRAK4* for experimental validation for five reasons: (i) *IRAK4* encodes interleukin 1 receptor-associated kinase 4 (IRAK4), a key mediator of signalling pathways upregulated in moPDAC-old and huPDAC-old tumours, including TLR and TNF signalling^44^. (ii) *IRAK4* was the only gene common to both prioritized groups (**Fig. 7A**; **Supplementary Table 14**). (iii) *IRAK4* was ‘exclusively’ located on the GN^moPDAC-old^, the GN^moMalign-old^, and the GN^huPDAC-old^ relative to corresponding younger GNs (**Supplementary Tables 5**, **7** and **11**). (iv) The CDS of *Irak4* ranked among the highest in both GN^moPDAC-old^ and GN^moMalign-old^ when compared with their respective young GNs (**Extended Data Fig. 7A-B**; **Supplementary Tables 5** and **11**). (v) Finally, although IRAK4 has previously been proposed as a therapeutic target in PDAC, the tumour contexts in which it may be most effective remain undefined^45–47^.

**Figure 7.**
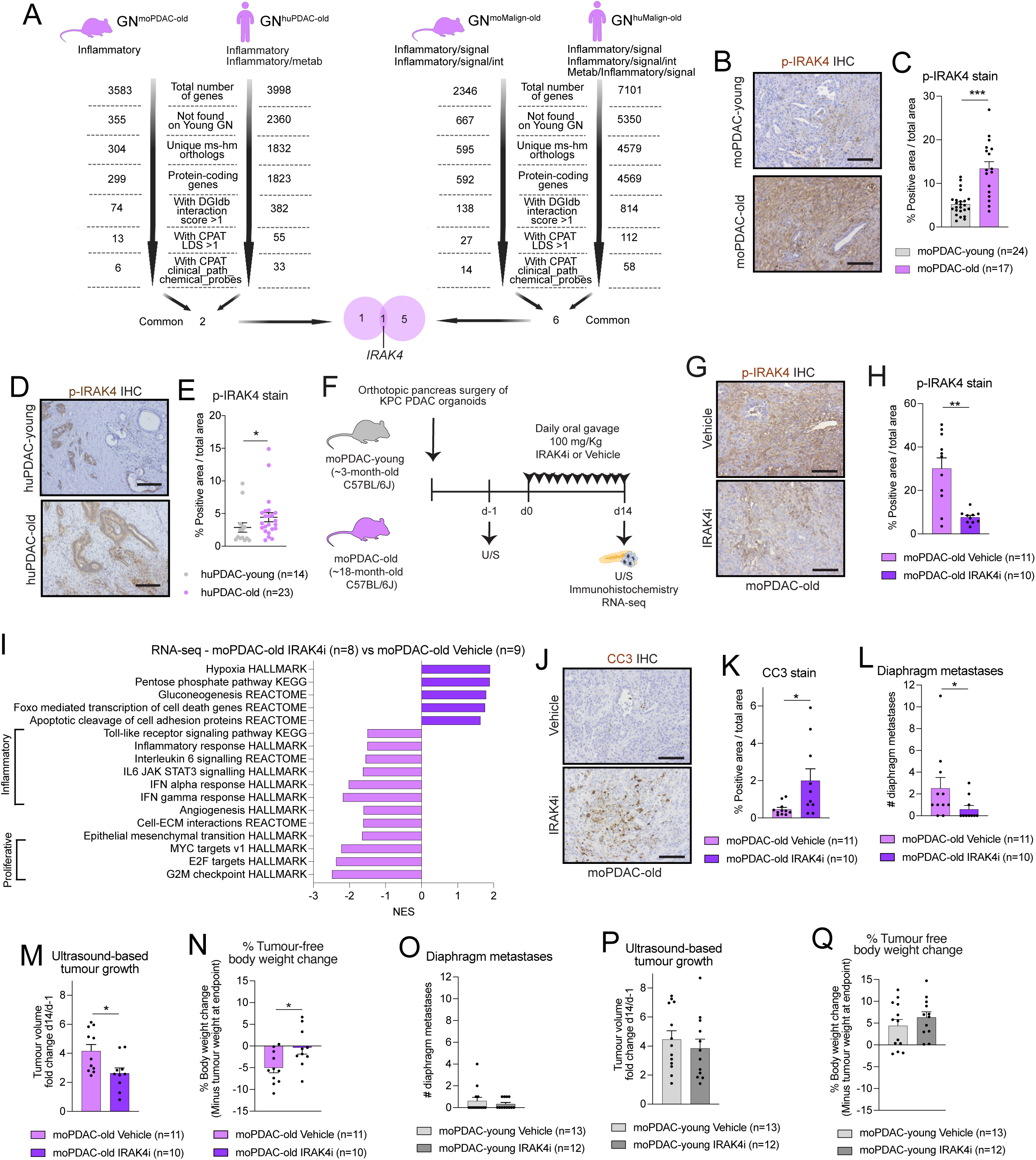
IRAK4 activity is a therapeutic vulnerability of old PDAC. **(A)** Left: Graphical representation of the prioritization strategy for candidate drug targets within the inflammatory communities on GN^moPDAC-old^ and GN^huPDAC-old^. Right: Graphical representation of the prioritization strategy for candidate drug targets within inflammatory-related communities on GN^moMalign-old^ and GN^huMalign-old^. *IRAK4* was the only common gene to both prioritization strategies. DGIdb, drug-gene interaction database. CPAT, canSAR.ai protein annotation tool. **(B)** Representative phospho-IRAK4 (p-IRAK4) stains in moPDAC-young and moPDAC-old. Scale bars, 50 μm. **(C)** Quantification of p-IRAK4 stain in moPDAC-young and moPDAC-old. Results show mean ± SEM. ***, *P* < 0.001, Mann-Whitney test. **(D)** Representative p-IRAK4 stains in huPDAC-young and huPDAC-old. Scale bars, 100 μm. **(E)** Quantification of p-IRAK4 stain in huPDAC-young and huPDAC-old. Results show mean ± SEM. *, *P* < 0.05, Mann-Whitney test. **(F)** Schematic of 2-week study in moPDAC-young and moPDAC-old tumour-bearing orthotopically grafted KPC PDAC organoid-derived mouse models with 100 mg/Kg IRAK4 inhibitor (IRAK4i, emavusertib, CA-4948) or vehicle by daily oral gavage. **(G)** Representative p-IRAK4 stains in vehicle- and IRAK4i- treated moPDAC-old. Scale bars, 50 μm. **(H)** Quantification of p-IRAK4 stain in vehicle- and IRAK4i- treated moPDAC-old. Results show mean ± SEM. **, *P* < 0.01, Mann-Whitney test. **(I)** Significantly upregulated and downregulated pathways identified by GSEA of IRAK4i-treated moPDAC-old (n=8) compared to vehicle-treated moPDAC-old (n=9). **(J)** Representative cleaved caspase 3 (CC3) stains in vehicle- and IRAK4i-treated moPDAC-old. Scale bars, 50 μm. **(K)** Quantification of CC3 stain in vehicle- and IRAK4i-treated moPDAC-old. Results show mean ± SEM. *, *P* < 0.05, Mann-Whitney test. **(L)** Number of diaphragm metastases of vehicle- and IRAK4i- treated moPDAC-old. Results show mean ± SEM. *, *P* < 0.05, Mann-Whitney test. **(M)** Tumour growth, as measured by ultrasound-based imaging, shown as ratio of tumour volumes at day 14 (d14) over tumour volumes at day -1 (d-1) of vehicle- and IRAK4i- moPDAC-old. Results show mean ± SEM. *, *P* < 0.05, Mann-Whitney test. **(N)** Body weight change in vehicle- and IRAK4i- moPDAC-old at day 14 compared to day -1. Results show mean ± SEM. *, *P* < 0.05, Mann-Whitney test. Body weight at endpoint was calculated by removing the tumour weight. **(O)** Number of diaphragm metastases of vehicle- and IRAK4i- treated moPDAC-young. Results show mean ± SEM. No significant difference was observed, as assessed by Mann-Whitney test. **(P)** Tumour growth, as measured by ultrasound-based imaging, shown as ratio of tumour volumes at day 14 (d14) over tumour volumes at day -1 (d-1) of vehicle- and IRAK4i- moPDAC-young. Results show mean ± SEM. No significant difference was observed, as assessed by Mann-Whitney test. **(Q)** Body weight change in vehicle- and IRAK4i- moPDAC-young at day 14 compared to day -1. Results show mean ± SEM. No significant difference was observed, as assessed by Mann-Whitney test. Body weight at endpoint was calculated by removing the tumour weight.

To determine whether IRAK4 activity differs across age groups, we compared total and phosphorylated IRAK4 (p-IRAK4) protein levels in young and old PDAC tumours from mice and humans. Remarkably, and in line with differential gene expression analyses, while total IRAK4 levels were similar across age groups, p-IRAK4 expression was significantly higher in PDAC-old compared to PDAC-young in both species (**Fig. 7B-E** and **Extended Data Fig. 7C-F**; **Supplementary Tables 5** and **7**). Thus, IRAK4 activity, not abundance, is elevated in aged PDAC, consistent with RECODR predictions. Interestingly, p-IRAK4 levels were also higher in young PDAC mouse models derived from KPPC (*Kras^LSL-G12D/+^*; *Trp53^fl/fl^*; *Pdx1*-*Cre*) organoids^29^, which carry homozygous *Trp53* deletion, compared with KPC organoid-derived moPDAC-young (carrying *Trp53^R172H^*), and were comparable to levels observed in old KPC and KPPC models (**Fig. 7C** and **Extended Data Fig. 7G-I**). These findings are in line with previous studies reporting efficacy of IRAK4 inhibition in ∼2-month-old mouse models of PDACs driven by deletion of either *Ink4a* or *Trp53*^45–47^.

Next, we randomised mice bearing KPC organoid-derived moPDAC-young or moPDAC-old tumours to receive the IRAK4 inhibitor emavusertib (IRAK4i, CA-4948^45^) or vehicle (**Fig. 7F**). In moPDAC-old, emavusertib significantly reduced p-IRAK4 levels and decreased pro-inflammatory pathways, including TLR, IFN and JAK-STAT signalling (**Fig. 7G-I**; **Supplementary Table 15**). Emavusertib also significantly downregulated genes associated with proliferation and epithelial-to-mesenchymal transition, while upregulating cell death programs in moPDAC-old tumours (**Fig. 7I**; **Supplementary Table 15**). These changes in gene expression correlated with increased levels of the apoptotic marker cleaved caspase 3 (CC3) in moPDAC-old tumours, reduced diaphragmatic metastasis burden, significant tumour growth inhibition, and attenuated body weight loss (**Fig. 7J-N**). In stark contrast, emavusertib had no significant effect on p-IRAK4 levels, TLR and JAK/STAT signalling, CC3 levels, tumour and metastatic burden, and body weight in moPDAC-young (**Fig. 7O-Q** and **Extended Data Fig. 7J-O**; **Supplementary Table 15**).

Altogether, our study shows that ageing profoundly remodels PDAC biology, creating age-specific therapeutic vulnerabilities. Specifically, our findings demonstrate that IRAK4 inhibition is selectively effective in old KRAS- and p53-mutant PDAC, highlighting the importance of incorporating age into preclinical model design and precision therapy development.

## DISCUSSION

Ageing is a major risk factor of PDAC, yet its impact on tumour biology and therapy response has been largely overlooked in preclinical studies. By combining organoid-derived PDAC models of different ages with cross-species graph-embedding analysis, we show that models in old mice better recapitulate the ageing-dependent biology of most PDAC patients (i.e., those aged ≥ 70 years) compared to those in younger mice. We also find that differences among mouse models can be leveraged to identify unique age-specific features and candidate therapeutic vulnerabilities of human PDAC. The use of a gene-centric, graph-based machine learning approach enabled by RECODR proved particularly effective in unravelling the complex biology of the heterogeneous cell populations that characterise PDAC at different ages. Particularly, we show that the aged TME drives a pro-inflammatory state that re-shapes PDAC malignant cell-stromal cell crosstalk and transcriptomes. These changes do not impact primary tumour growth but instead alter therapy response, underscoring the need for age-informed treatment strategies. This is relevant to the human disease, as evidence suggests that tailored treatments may be required for frailer older patients despite no evidence of faster PDAC growth kinetics in this patient group^6,48^.

Chronic inflammation is a hallmark of ageing in health and disease^47^. For example, the aged murine pancreas has been found to be associated with chronic inflammation and stromal activation in response to injury^49^. Yet, age-associated inflammation has not been previously described in PDAC. Here, we find that ageing increases inflammatory programs in PDAC tumours, including IRAK4 activity, creating age-specific therapeutic vulnerabilities. Further work will be required to understand how this pro-inflammatory phenotype emerges and is maintained through immune-stromal-malignant cell cross-talks. Such understanding will help identify additional therapeutic targets.

Among the age-dependent TME changes observed in our models, macrophage abundance decreased in moPDAC-old. This is consistent with prior studies that showed reduced macrophage numbers, while upregulating pro-inflammatory programs, in skin wounds in old (22-24-month-old) relative to young (2-3-month-old) mice^50^. Aged macrophages were also shown to have reduced phagocytosis, chemotaxis and migration capacity relative to those in normal human and mouse young tissues^51^. Conversely, MHCII^+^ macrophages, which are known to be associated with M1-like activation^52^, were enriched in moPDAC-old, aligning with RNA-seq deconvolution studies that reported enrichment of M1 macrophages in older human PDAC^53^. In addition to macrophages, neutrophil and fibroblast populations were also markedly shaped by ageing in moPDAC-old, indicating that ageing broadly remodels stromal composition. These changes should be considered when seeking to understand PDAC age-dependent biology and tailor therapies.

Prior studies have also pinpointed cellular senescence as a hallmark of ageing and tumorigenesis^54^. This has included the emergence of immunosuppressive p16^+^ senescent myCAF subsets in PDAC and breast cancer^55–57^. While our analysis did not identify expansion of this senescent CAF population in moPDAC-old, further research will be required to identify the impact of senescence on the TME, tumour progression and therapy response in aged PDAC.

As done for other cancer types^21–24^, our study leveraged transplantation-based mouse models to determine the impact of differently aged stroma on malignant cells generated from a common source. Extending this framework to models in which both stromal and epithelial compartments are chronologically old, an approach that has so far been explored only in lung cancer^58^, will further refine our understanding of age-dependent malignant-stroma interactions. Addressing this question will require the use of inducible GEMMs that enable effective PDAC formation in aged mice. Future studies should also investigate how ageing interacts with other malignant cell-intrinsic factors, including tumour stage, sex and distinct mutational signatures.

Most clinical trials in PDAC have been unsuccessful and survival rates have improved only marginally over the last twenty-five years^59^. The pressure for more effective treatments has driven the conduct of clinical trials often with only limited preclinical evidence to support them^60^. Models that more faithfully recapitulate specific types of PDAC may help design more precise and effective treatment strategies. Previous studies of the impact of the ageing TME on PDAC growth using 2D cell-derived models have yielded conflicting results^26,27,61^. Here, by leveraging organoid-derived PDAC models, and consistent with clinical observations, we find that TME age does not accelerate PDAC growth kinetics but instead reshapes therapeutic vulnerability states.

We focused on discovering vulnerabilities in old PDAC since this age group constitutes most patients at diagnosis, is less likely to receive adjuvant therapy, and shows the worse outcomes after tumour resection^6,7^. While clinical outcomes in older patients are influenced by frailty^48^, our findings demonstrate that ageing also imposes distinct, targetable biological states within PDAC tumours. The biological insights provided by our analyses may lead to age-specific biomarkers and associated treatments to guide precision therapeutic strategies for distinct PDAC groups. Such an approach will demand better understanding of intermediate PDAC, which has been largely overlooked as a unique age group relative to young and old PDAC patients^33,62,63^. Indeed, our analyses suggest that huPDAC-intermediate has unique biological features that are distinct from tumours diagnosed in young or old patients.

As a first step to identify context-specific therapeutic vulnerabilities to molecular targeted therapies in PDAC, we tested and validated the RECODR prediction that inhibition of IRAK4 would be effective in old but not young moPDAC models. IRAK4 activity was upregulated in moPDAC-old, despite similar expression levels, and its inhibition impaired tumour growth only in aged models. Since moPDAC-old tumour growth kinetics are comparable to moPDAC-young tumours, these data suggest that alternative tumour-promoting programs are active in moPDAC-young tumours. Previous studies have indicated IRAK4 as a therapeutic target in a variety of malignancies, including PDAC^45–47,64–66^, although the precise context in which this might be most effective was not defined. Here, we show that IRAK4 inhibition does not impair the growth or metastasis of KRAS^G12D^, p53-mutant organoid-derived PDACs orthotopically transplanted in ∼3-month-old mice but is effective against these PDACs in ∼18-month-old mice. Notably, we also find that p-IRAK4 levels were higher in young PDAC mouse models driven by p53 deletion compared to p53-mutant moPDAC-young models, and were not further modulated by ageing in this context. Thus, IRAK4 activity, and age-specific modulation, varies in response to PDAC genetic context. Importantly, our models are driven by p53 mutation, which is far more common than p53 deletion in human PDAC^67^. Our findings are also consistent with prior studies that have shown that distinct genetics, including p53 mutation versus deletion, differently impact PDAC therapy response^29,35,68–70^. Most importantly, our study highlights the context-dependent nature of therapeutic targets and the critical role of the aged TME in shaping PDAC biology and treatment response.

## MATERIAL AND METHODS

### Human PDAC tissues

Human PDAC tissues used in this study were obtained from surgical resections performed at Addenbrooke’s hospital (Cambridge, UK). Written informed consent was acquired from patients before specimens’ acquisition, and formalin-fixed paraffin-embedded (FFPE) specimens were bio-banked by a pathologist at the Human Research Tissue Bank. Specimens were acquired using the CAMPAN (REC 08/H0306/32), Pancrea_tive-T (REC 22/EE/0233) or Pancrea_tive (REC21/WM/0119) protocols. All samples were anonymized to the researchers. Available clinical information of the human PDAC tissues used for histological analyses is provided in **Supplementary Table 1**. Age groups were defined considering previous literature on early-onset (≤ 55 years) and average-age onset (≥ 70 years) PDAC^33,63^.

### Mouse models

Males and females young, intermediate and old C57BL/6J mice were purchased from the Charles River Laboratory (strain number 632) or from Janvier Laboratories (C57BL/6JRj; RRID:MGI:2670020). At the Cancer Research UK Cambridge Institute (CRUK-CI), all animals were housed in accordance with the guidelines of the UK Home Office “Code of Practice for the Housing and Care of Experimental Animals”. They were kept behind strict barriered housing, which maintains animals at a well-defined microbiological health status. All animals were fed expanded rodent diet (Labdiet) and filtered water *ad libitum*. Environmental enrichment included nesting material, chew blocks, and an area to retreat. All animal procedures and studies were reviewed by the CRUK-CI AWERB, approved by the Home Office and conducted under PPL number PP4778090 in accordance with relevant institutional and UK national guidelines and regulations.

### Orthotopic pancreas transplantation mouse models

Orthotopic injections were conducted as previously described^71^. Briefly, 10,000 single cells prepared from organoid cultures were resuspended as a 35 μL suspension of 50% Matrigel (354230; Corning) in PBS and injected into the pancreas of ∼8-15-week-old (∼2-3-month-old) mice, ∼51-55-week-old (∼12-13-month-old) mice or ∼77-87-week-old (∼18-20-month-old) male and female C57BL/6J mice. Pancreatic tumours were imaged using the Vevo 2100 Ultrasound at two different orientations with respect to the transducer. Tumour volumes were typically measured at two angles using the Vevo LAB software program (version 5.7.0). Only mice with successful (i.e., non-leaked/re-injected) orthotopic injections were included in tumour volume and metastasis analyses. Tumour volume analyses were performed blindly prior to plotting the data for visualization.

### *In vivo* IRAK4 inhibition study

The IRAK4 inhibitor emavusertib was prepared daily as a suspension in 0.25% w/v hydroxyethyl-cellulose (09368; Sigma-Aldrich) and 0.5% w/v Tween-20 (P1379; Sigma-Aldrich) in sterile water and sonicated for five minutes. Mice were administered vehicle or 100 mg/Kg of emavusertib (CA-4948; HY-135317, MedChem Express) for 14 days, once a day (in the AM) via oral gavage^45,72^. Pancreatic tumours were imaged prior to enrolment (day -1) and at endpoint (day 14) using the Vevo 2100 Ultrasound at two different orientations with respect to the transducer. Mice with ∼5-8 mm tumour diameters were randomized and enrolled one day after scanning. Tumour volumes were typically measured at two angles using the Vevo LAB software program (version 5.7.0), and growth rate was calculated by dividing the volume at day 14 for the volume at day -1. Tumour volume analyses were performed blindly prior to plotting the data for visualization.

### Organoid lines and culture

Murine PDAC KPC (*Kras^LSL-G12D/+^*; *Trp53^LSL-R172H/+^*; *Pdx1*-*Cre*) organoid lines T69A (female) and T6-LOH (male), and the murine PDAC KPPC (*Kras^LSL-G12D/+^*; *Trp53^fl/fl^*; *Pdx1*-*Cre*) organoid line T113, were previously published and maintained as previously described^29,73,74^. Organoids were typically cultured for no more than 30 passages at 37°C with 5% CO2.

### Immunohistochemical and histological analyses

Standard procedures were used for immunohistochemistry. Primary antibodies were anti-mouse αSMA (ab5694; Abcam; RRID:AB_2223021), anti-human αSMA (M0851, DAKO; RRID:AB_2223500), CC3 (9664; Cell signaling technologies; RRID:AB_2070042), IRAK4 (4363; Cell Signaling Technologies; RRID:AB 2126429), and p-IRAK4 (MAB2538; Abnova; RRID:AB_10555313). Hematoxylin (H-3404; Vector Lab) was used as nuclear counterstain. Hematoxylin and eosin stains and Masson’s trichrome stains were performed according to standard protocols by the Histopathology core at the CRUK-CI. Brightfield images of tissue slides were obtained with an Axio Vert.A1 (ZEISS). Stained sections were scanned with Aperio ScanScope CS and analyzed using the ImageScope Positive Pixel Count algorithm. For Masson’s trichrome stains, the percentage of collagen area was determined by calculating the percentage of blue pixels relative to the entire stained area. To quantify αSMA, CC3, IRAK4, and p-IRAK4 stains, the percentage of positive pixels was calculated relative to the entire tissue section. For Masson’s trichrome, p-IRAK4, IRAK4 and αSMA quantifications of human PDAC tissues, only the PDAC area was analysed following annotations from a pathologist (R.B.). Quantifications were performed blindly prior to plotting the data for visualization.

### Flow cytometry

PDAC tumours from male and female mice were processed as previously described^35,36^. Briefly, cells were blocked for 15 minutes on ice with CD16/CD32 Pure 2.4G2 (553142, BD Bioscience). For flow cytometric analysis of CAFs, CD90^+^ CAFs, endothelial cells, epithelial cells and immune cells, cells were stained for 30 minutes on ice with anti-mouse CD31-PE/Cy7 (102418; BioLegend; ; RRID:AB_830757), CD45-PerCP/Cy5.5 (103132; BioLegend; RRID:AB_893344), CD326 (EpCAM)-AlexaFluor 488 (118210; BioLegend; RRID:AB_1134099), PDPN-APC/Cy7 (127418; BioLegend: RRID:AB_2629804), MHCII-BV785 (107645; BioLegend; RRID:AB_2565977), Ly6C-APC (128015; BioLegend; RRID:AB_1732087), and CD90-PE (ab24904; Abcam; RRID:AB_448474).

For flow cytometric analysis of CD49E^+^ CAFs and CD56^+^ CAFs, cells were stained for 30 minutes on ice with CD326 (EpCAM)-AlexaFluor 488 (118210; BioLegend; RRID:AB_1134099), CD31-AlexaFluor 488 (102413; BioLegend; RRID:AB_493409), CD45-AlexaFluor 488 (103122; BioLegend; RRID:AB_493531), PDPN-APC/Cy7 (127418; BioLegend; RRID:AB_2629804), CD49e-BV76 (740863; BioLegend; RRID:AB_2870652), CD56-APC (FAB7820A; Biotechne; RRID:AB_3652722), SCA1-PE (108107; Biolegend; RRID:AB_313344), CD26-PerCP/Cy5.5 (45-0261-80; Thermo Fisher Scientific; RRID:AB_1548741), CD73-PE/Cy7 (127223; Biolegend; RRID:AB_2716102).

For flow cytometric analysis of CD105^+^ CAFs, cells were stained for 30 minutes on ice with CD326 (EpCAM)-AlexaFluor 488 (118210; BioLegend; RRID:AB_1134099), CD31-AlexaFluor 488 (102413; BioLegend; RRID:AB_493409), CD45 AlexaFluor 488 (103122; BioLegend; RRID:AB_493531), PDPN-APC/Cy7 (127418; BioLegend; RRID:AB_2629804), CD105-PE/Cy7 (120409; Biolegend; RRID:AB_1027702), CD200-PE (123807; Biolegend; RRID:AB_2275651), MHCII-BV785 (107645; BioLegend; RRID:AB_2565977), CD55-APC (131811; Biolegend; RRID:AB_2800631), and CD106-PerCP/Cy5.5 (105716; Biolegend; RRID:AB_1595489).

For flow cytometric analysis of T cells, B cells and NK cells, cells were stained for 30 minutes on ice with anti-mouse CD45-PerCP/Cy5.5 (103132; BioLegend; RRID:AB_893344), TCR-β-Alexa488 (109215; BioLegend; RRID:AB_493344), CD3e-Alexa488 (100321; BioLegend; RRID:AB_389300), CD8-APC/Cy7 (100713; BioLegend; RRID:AB_312752), CD4-APC (100515; BioLegend; RRID:AB_312719), CD19-PE/Cy7 (115520; Biolegend; RRID:AB_313655), NK1.1-BV785 (108749; Biolegend; RRID:AB_2564304), and TIGIT-PE (142103; BioLegend; RRID:AB_10895760).

For flow-cytometric analysis of macrophages, neutrophils and dendritic cells, cells were stained for 30 minutes on ice with anti-mouse CD45-PerCP/Cy5.5 (103132; BioLegend; RRID:AB_893344), CD11b-PE/Cy7 (101215; BioLegend; RRID:AB_312798), F4/80-BV785 (123141; BioLegend; RRID:AB_2563667), MHCII-APC/Cy7 (107627; BioLegend; RRID:AB_1659252), CD11c-APC (117309; BioLegend; RRID:AB_313779), Gr1-PE (108407; BioLegend; RRID:AB_313372).

After staining, cells were resuspended in PBS with DAPI and analysed on a BD FACSymphony cell analyser. Flow analyses were performed blindly prior to plotting the data for visualization.

### Single-cell RNA-sequencing data analyses of murine PDAC tumours

scRNA-seq data of moPDAC tumours are available at the Gene Expression Omnibus (GEO) under the accession number GSE263595. scRNA-seq analyses of moPDAC tumours are included in **Supplementary Tables 2-4**. PDAC tumours from male and female mice were processed as previously described^35,36^. Single cells were submitted in 55 μl at a concentration of 370 cells per μl and were processed by the CRUK-CI Genomics core with 10x Genomics 3’V3.1 chemistry and sequenced on NovaSeq6000 (Illumina) or NovaSeqX (Illumina). Cell Ranger (10x Genomics) workflow^75^ was used to align FASTQ files to the mm10 (GENCODE vM23/Ensembl98) version 2020-A (Jul 7, 2020) mouse transcriptome reference genome that was downloaded from the 10x Genomics database to generate the raw counts of gene expression quantities. From the raw counts of each sample, ambient RNA removal was performed using CellBender with false rate threshold of 0.01^76^. Downstream analysis was performed using Scanpy^77^. Cells expressing less than 200 genes, genes detected in less than 3 cells, cells with more than 5% of mitochondrial gene content, and cells with total UMI counts greater than 40000 were discarded. Doublets were detected using SOLO, as implemented in scvi-tools^78^ on each sample separately. Then, filtered samples were concatenated into a single AnnData object. Raw counts were normalized to a total of 10000 reads per cell and log-transformed. Highly variable genes were identified using “Seurat_v3” as a flavor method, and the top 4000 genes were selected for dimensional reduction. Principal component analysis (PCA) was performed, the top 30 PCs were used to construct the k-nearest-neighbour graph, and to perform unsupervised clustering using the Leiden algorithm. Uniform manifold approximation and projection (UMAP) plots were used to visualize the datasets in 2-dimentional space. Cell clusters were annotated based on specific markers (**Extended Data Fig. 1K**).

#### Copy number variation (CNV) analysis

A python implementation of InferCNV of the Trinity CTAT Project (https://github.com/broadinstitute/inferCNV) was used to estimate the copy number status in each cell type. Fibroblasts as reference key and a 250-genes window size were used (**Extended Data Fig. 1L**).

#### Pseudotime and differentiation potential analysis

Differentiation potential was performed using CellRank tool on malignant cell and neutrophil clusters^79^. The Cytotrace kernel was used to estimate cellular differentiation states based on transcriptional diversity^80^. Following the official tutorial, the normalized counts in adata.X were set to both spliced and unspliced layers of the AnnData object. Then, Cytotrace scores were computed using the compute_cytotrace function and ct_pseudotime score was used to visualize the relative differentiation potential across cells.

#### Differential expression analysis

The AnnData object was converted to Seurat object. Then, the Model-based Analysis of Single-cell Transcriptomics (MAST)^81^ R package was used for differential expression analysis. MAST implementation in Seurat FindMarkers function was used with the following parameters: min.pct = 0.05 and logfc.threshold = 0 to define the differentially expressed genes (DEGs) between conditions (e.g. old versus young) in each cell cluster. Then, the genes were pre-ranked according to their avg_log2FC as a prior step for GSEA. Thereafter, GSEA was performed using the clusterprofiler R package against pathways from Hallmark, Reactome, and C2 canonical pathway collection (C2.cp.v5.1) downloaded from the Molecular Signatures Database (MSigDB)^82,83^. GSEA significance was defined as NES >1.5 or <-1.5 and FDR < 0.25 as defined by Liberzon et al^84^.

### Single-cell RNA-sequencing analyses of KPC GEMMs

KPC GEMM scRNA-seq data are from Elyada *et al* ^34^. The dataset is publicly available at the GEO under accession number GSE129455 (**Supplementary Table 9**). Raw sequencing data were downloaded using the sra-tools (https://github.com/ncbi/sra-tools). Raw data preprocessing and downstream analyses were performed using the same pipeline described in the method section above.

### Single-cell RNA-sequencing analyses of human PDAC tumours

The raw count matrix in “pk_all.rds” from Chijimatsu et al. (2022)^40^ was utilized. Samples with “Tumour” and available age annotation were selected, resulting in a total of 55 samples from the CA001063^85^, GSE154778^86^, GSE155698^87^, and OUGS^40^ projects (**Supplementary Table 6**). The raw counts were processed using the Scanpy package^77^. Cells expressing less than 200 genes and genes that are detected in less than 3 cells were filtered out. Counts were normalized and log-transformed using scanpy function sc.pp.normalize_total with target sum 10000 and sc.pp.log1p. Thereafter, highly variable genes were identified using “seuratv3” as a flavor method in Scanpy highly variable gene function, and the top 4000 genes were selected for PCA.

Top 30 PCs were used to construct k-nearest-neighbor graph, compute UMAP embedding, and to perform clustering with the Leiden algorithm. Potential batch effect and confounding factors were systematically evaluated. Data integration was performed in the reduced PCA space using Harmony python implementation, with “Project ID” and “Sample ID” as batch covariates^88^. Integration quality was assessed using the adjusted mutual information^88^ (AMI) (**Supplementary Table 6**), which quantifies the agreement between cluster assignment and identity labels. The AMI score was computed using the adjusted_mutual_info_score function in the scikit-learn package. Clustering was performed using the Leiden algorithm^89^. All cell clusters were annotated based on specific cell marker genes (**Extended Data Fig. 4C**) and malignant cell cluster was validated through CNV scoring across all cell clusters (**Extended Data Fig. 4D**) using the Python implementation of InferCNV as described above. Age groups were defined considering previous literature on early-onset (≤ 55-year-old) and average-age onset (≥ 70-year-old) PDAC^33,63^.

### RECODR analyses

To generate and analyse gene co-expression GNs using scRNA-seq datasets of PDAC tumours from different age groups, we used the previously published RECODR pipeline^39^. RECODR analyses of murine and human PDAC are included in **Supplementary Tables 5**, **9** and **11**, and **Supplementary Tables 7** and **12**, respectively. Initially, the processed AnnData was first split by age group. For each age group, log-normalized expression counts were converted into dataframes by using Scanpy function to_df(), with cells as columns and genes as rows. These dataframes were then saved as csv within the same directory to be used as input for RECODR. RECODR then performs the following steps: first, the count tables were subset to include genes that were expressed in at least 5% of cells. Next, a pairwise Spearman correlation analysis was carried out between every set of genes in the subset counts for each age. The correlations were then subset to include gene pairs that were correlated equal to or above a Spearman correlation of 0.2, adjusted Bonferroni P-value of less than 0.05 and self-correlations removed. Next, undirected and unweighted graph networks were made with the Python package NetworkX using the from_pandas_edgelist function, with nodes as genes and edges connecting two nodes if they were correlated equal to or above a Spearman correlation of 0.2. Community detection of each graph was run with the best_partition function of the Python-louvain library. Next the Python package Pecanpy was used for an optimized version of the Node2Vec model^90^. First, NetworkX graph edges for each graph were converted to a format suitable for Pecanpy using the write_edgelist function from NetworkX with the ‘delimiter’ argument set to “ ‘ ‘ “ and ‘data’ argument set to ‘False’. Graphs in the format appropriate for Pecanpy were made using the read_edg function with ‘weighted’ and ‘directed’ set to False, and ‘delimiter’ set to “ ’ ’ ". Next, graph walks were run in Pecanpy with the ‘p’ parameter set to ‘1’ and the ‘q’ parameter set to ‘0.7’. Next, the simulate_walks function was used to generate graph walks with ‘num_walks’ set to 200 and ‘walk_length’ set to 80. The walks were then fed directly to the Word2Vec model^91^ from natural language processing library Gensim with a ‘window’ size set to 10, ’sg’ set to 1 (to ensure the skip-gram objective was used), ‘vector_size’ set to 100 and ‘epochs’ set to 25. All other Word2Vec arguments were kept to their default parameters. After training, the Procrustes alignment as described in Jassim et al^39^ was run to align word embeddings of common genes between different ages and the cosine similarity taken between vectors of the same gene across different ages to approximate context drift. RECODR was run on High computing cluster (HPC) with cpu = 38. Only communities with 35 genes or more were included in downstream analyses. Functional annotation of each community was performed using their gene list and a pathway enrichment analysis was performed using the clusterprofiler R package against the Hallmark, Reactome, and C2 canonical pathway collection (C2.cp.v5.1) downloaded from the MSigDB^82,83^.

#### Gene context drift score

Cosine similarity between vectors was calculated as one minus the cosine distance, yielding values between -1 and 1. The CDS for each gene was then derived using RECODR’s default parameters based on these similarity values. To further assess network-level changes, we computed a NodeScore for each gene across all pairwise age group comparisons (old vs young, intermediate vs young, old vs intermediate) using the NodeScore function in RECODR^39^. Briefly, for each gene (index gene), the first score is the context drift of the index gene relative to itself in the comparator graph; the second score takes the average of the context drift of genes connected to the index gene as an approximation to the amount of transcriptional remodelling happening around the index gene; the third score is the hypothesized reachability of a gene on the graph network calculated as the number of unique connections to an index gene plus the number of unique connections to those genes; the fourth score is the unique number of genes connected to an index gene that were new to that graph and not present in the comparator graph. The sum of all four scores was taken as the final CDS score. The resulting output is a CSV file listing genes alongside their relative neighbourhood scores and drift scores.

#### GN representation

The "Networkx" Python library was used to visualize gene networks^92^. The connections (edges) between genes were determined based on co-expression scores (Spearman correlation >= 0.2 and FDR < 0.05). The graph gpickle files from RECODR output were used for plotting, kamada_kawai_layout^93^ from networkx was used for the node position and the size of each node indicated by using the nodes betweenness scaled to 1000. Specifically, the betweenness of each gene was calculated using the "centrality.betweenness_centrality" function from the "networkx.algorithms" module. Betweenness centrality is a metric that quantifies how much a node controls the flow of information within a network: this value ranges from 0 to 1 and was scaled by a factor of 5000 to enhance visual representation. Small communities (i.e., < 35 genes present) are included in the GNs (in light grey) but were neither further analysed nor included in other plots.

#### DEG versus CDS plots

To compare CDS with DEG profiles, we used the final CDS generated by RECODR for each gene. Differential expression analysis (DEA) was performed using the MAST method, implemented via the FindMarkers function in Seurat with with logfoldchange cutoff = 0 and min_pct = 0.05^94^. This allowed for a direct comparison between genes exhibiting context drift and those identified as differentially expressed across age groups. DEA analysis of immune cells was performed on the combined macrophages, dendritic cells and neutrophils cell clusters.

#### Mouse to human gene orthologs

To obtain human orthologs from mouse gene symbols, we downloaded the ‘’mouse/human orthology with phenotype annotations’’ list from the Mouse Genomic informatics portal (https://www.informatics.jax.org). This list is composed by 28,880 mouse genes with available human orthologs. To obtain unique mouse to human orthologs, we eliminated redundant mouse to human orthologs (i.e., mouse or human genes that have multiple corresponding genes in the mouse or human genome, respectively). After the filtering, the final list is composed by 17,094 non-redundant, mouse to human orthologs including *CDKN2A* and *CDKN2B* (**Supplementary Table 8**).

#### Candidate target genes prioritization strategy

To find genes that were candidate therapeutic vulnerabilities of old murine and human PDAC, we first selected genes present across the inflammatory-related communities of GN^moPDAC-old^ (O1 inflammatory), GN^moMalign-old^ (O4 inflammatory/cell signalling and O5 inflammatory/cell signalling/interactions), GN^huPDAC-old^ (O2 inflammatory and O4 inflammatory/metabolism) and GN^huMalign-old^ (O2 inflammatory/cell signalling, O3 metabolism/inflammatory/cell signalling and O4 inflammatory/cell signalling/interactions). We selected genes that were ‘exclusive’ to GNs of old, but not young, PDAC. We then converted the genes into unique human orthologs using the list of 17,094 non-redundant mouse to human orthologs mentioned above. We selected genes from the communities listed above that have a gene-drug pair and an interaction score > 1 based on the drug-gene interaction database (DGIdb) (www.dgidb.org)^42^. From this pool, we then selected genes that have a ligand druggability score (LDS) > 1 and validated chemical probes (i.e., compounds that have evidence for their selectivity and potency against the protein product of a given gene) based on the canSAR.ai protein annotation tool (CPAT) (www.cansar.ai/cpat)^43^. Finally, to obtain common targets across comparisons, we overlapped prioritized genes between GN^moPDAC-old^ and GN^huPDAC-old^ (i.e., all cell analyses) and between GN^moMalign-old^ and GN^huMalign-old^ (i.e., malignant cell-only analyses) (**Fig. 7A**; **Supplementary Table 14**).

### RNA-sequencing of IRAK4i-treated PDAC tumours

RNA-seq data of vehicle- and IRAK4i- treated murine tumours are available at the GEO under the accession number GSE302662. Bulk RNA-seq analyses of vehicle- and IRAK4i- treated PDAC are included in **Supplementary Table 15** and were performed by the CRUK-CI Bioinformatics core. PDAC tumour samples from male and female mice were snap-frozen and stored at -80 °C. Tumour samples were thawed and dissociated in 0.8 mL of TRIzol Reagent using steel beads in a Qiagen TissueLyser II (85300; Qiagen) for 4 minutes at 25 Hz. RNA was extracted using the PureLink RNA mini kit (12183018A; Invitrogen). RNA concentration was measured using a Qubit and RNA quality was assessed on a TapeStation 4200 (Agilent) using the Agilent RNA ScreenTape kit. mRNA library preparations were performed by the CRUK-CI Genomics core using 55 μL of 1000 ng of RNA per sample (RNA integrity number > 6). Illumina libraries were then sequenced on 2 lanes of SP flowcell on NovaSeq6000 (Illumina). FASTQ files were aligned, and the expression levels of each transcript were quantified using Salmon (v1.4.0)^95^ with the annotation from ENSEMBL (GRCh38 release 98) with recommended settings. Transcript-level expression was loaded and summarized to the gene level by using tximport^96^. Differential gene expression was performed using DESeq2 (v2)^97^ by applying lfcshrink function^98^. The principal components for variance-stabilized data were estimated using plotPCA function, available in DESeq, and ggplot2 (https://ggplot2.tidyverse.org). Following differential gene expression, genes were pre-ranked based on the negative logarithmic p-value and the sign of the Log2 fold change. GSEA was performed using clusterprofiler against the Hallmark, Reactome, and C2 canonical pathway collection (C2.cp.v5.1) downloaded from the MSigDB^82,83^. GSEA significance was defined as NES >1.5 or <-1.5 and FDR < 0.25 as defined by Liberzon et al^84^.

### Statistical analysis and graphical representations

GraphPad Prism software, customized R scripts and Jupyter notebooks were used for statistical analysis, simple linear regressions, and graphical representation of data. Alluvial plots were generated using the open-source software RAWGraphs 2.0^99^. Mouse and human icons were downloaded from www.bioicons.com. Statistical analysis was performed using non-parametric Mann-Whitney U test, Kruskal-Wallis test followed by Dunn’s post-hoc pairwise comparisons with Benjamini-Hochberg FDR correction, hypergeometric test (using 17,094 orthologs as the background set), F-test, Fisher’s test, and chi-square test, as appropriate. Adjustment for multiple comparisons was applied where relevant. Covariates were evaluated when appropriate including batch correction of RNA-sequencing analysis. Statistics on proportional composition graphs for scRNA-seq data were calculated using scCODA^100^. All statistical parameters, including the definition of significance thresholds, number of replicates, and exact tests used, are reported in the corresponding figure legends and/or figure panels.

## ACKNOWLEDGEMENTS

The authors would like to thank the CRUK-CI Biological Resources Unit, Genomics, Bioinformatics, Flow Cytometry, Imaging, Compliance and Biobanking, Scientific Computing, and Histopathology core facilities. The authors would also like to acknowledge the Human Tissue Research Bank, which is supported by the NIHR Cambridge Biomedical Research Centre (NIHR203312). This work was mainly supported by a Cancer Research UK (CRUK) institutional grant (A27463), which also supported J.S.M., S.M., D.M., M.Z., and S.P.T., and a UKRI Future Leaders Fellowship of which G.B. is recipient and that also supported W.L and E.G.L. G.B. also received a Pancreatic Cancer Research Fund grant and a US Department of Defense PCARP grant, which supported G.M., M.V. and J.S.M., a NCI-CRUK Cancer Grand Challenge grant, which supported M.J., W.K.L. and S.H., and a Pancreatic Cancer UK Future Leaders Academy grant, which supported P.S.W.C. J.A.H. was supported by a Harding Distinguished Postgraduate Program PhD studentship (Cambridge Trust). E.G.L. was initially supported by an MRC Doctoral Training Grant. The authors also want to thank Pancreatic Cancer UK that supported G.M. through a Career Foundation Fellowship award. Finally, we acknowledge initial funding support from the Royal Society and the Isaac Newton/Wellcome ISSF/University of Cambridge scheme.

## AUTHOR CONTRIBUTIONS

J.A.H and M.J. designed and conducted the experiments, and revised the paper. G.M., E.G.L, W.L., D.M., M.Z., and A.J. conducted the experiments, and revised the paper. S.P.T, J.S.M., W.K.L, S.H., P.S.W.C. and S.M. conducted the experiments. R.B. performed histopathology analysis of human tissues. M.V. co-supervised the mouse studies. R.J.G. co-supervised the RECODR analyses and revised the paper. G.B conceptualised and supervised all studies, designed and conducted the experiments, and wrote the paper.

## REFERENCES

1. Siegel, R. L., Kratzer, T. B., Giaquinto, A. N., Sung, H. & Jemal, A. Cancer statistics, 2025. CA Cancer J Clin 75, 10–45 (2025).

2. Biffi, G. & Tuveson, D. A. Diversity and Biology of Cancer-Associated Fibroblasts. Physiol Rev 101, 147–176 (2021).

3. Falcomatà, C. et al. Context-Specific Determinants of the Immunosuppressive Tumor Microenvironment in Pancreatic Cancer. Cancer Discov 13, 278–297 (2023).

4. Ma, R.-Y., Black, A. & Qian, B.-Z. Macrophage diversity in cancer revisited in the era of single-cell omics. Trends Immunol 43, 546–563 (2022).

5. Ng, M. S. F. et al. Deterministic reprogramming of neutrophils within tumors. Science 383, eadf6493 (2024).

6. GBD 2017 Pancreatic Cancer Collaborators. The global, regional, and national burden of pancreatic cancer and its attributable risk factors in 195 countries and territories, 1990-2017: a systematic analysis for the Global Burden of Disease Study 2017. Lancet Gastroenterol Hepatol 4, 934–947 (2019).

7. He, J. et al. Young patients undergoing resection of pancreatic cancer fare better than their older counterparts. J Gastrointest Surg 17, 339–344 (2013).

8. Ansari, D. et al. Relationship between tumour size and outcome in pancreatic ductal adenocarcinoma. Br J Surg 104, 600–607 (2017).

9. Alese, O. B. et al. Young Adults With Pancreatic Cancer: National Trends in Treatment and Outcomes. Pancreas 49, 341–354 (2020).

10. Beeghly-Fadiel, A. et al. Early onset pancreatic malignancies: Clinical characteristics and survival associations. Int J Cancer 139, 2169–2177 (2016).

11. Ordonez, J. E. et al. Clinicopathologic Features and Outcomes of Early-Onset Pancreatic Adenocarcinoma in the United States. Ann Surg Oncol 27, 1997–2006 (2020).

12. Piciucchi, M. et al. Early onset pancreatic cancer: risk factors, presentation and outcome. Pancreatology 15, 151–155 (2015).

13. Ramai, D. et al. Early- and late-onset pancreatic adenocarcinoma: A population-based comparative study. Pancreatology 21, 124–129 (2021).

14. Tas, F., Sen, F., Keskin, S., Kilic, L. & Yildiz, I. Prognostic factors in metastatic pancreatic cancer: Older patients are associated with reduced overall survival. Mol Clin Oncol 1, 788–792 (2013).

15. He, W. et al. Underuse of surgical resection among elderly patients with early-stage pancreatic cancer. Surgery 158, 1226–1234 (2015).

16. Ansari, D., Althini, C., Ohlsson, H. & Andersson, R. Early-onset pancreatic cancer: a population-based study using the SEER registry. Langenbecks Arch Surg 404, 565–571 (2019).

17. Henry, C. J. & DeGregori, J. Modelling the ageing dependence of cancer evolutionary trajectories. Nat Rev Cancer 1–24 (2025) doi:10.1038/s41568-025-00838-3.

18. Lloyd, E. G., Henríquez, J. A. & Biffi, G. Modelling the micro- and macro- environment of pancreatic cancer: from patients to pre-clinical models and back. Disease Models & Mechanisms 17, dmm050624 (2024).

19. Wang, S., Lai, X., Deng, Y. & Song, Y. Correlation between mouse age and human age in anti-tumor research: Significance and method establishment. Life Sciences 242, 117242 (2020).

20. Fane, M. & Weeraratna, A. T. How the ageing microenvironment influences tumour progression. Nat Rev Cancer 20, 89–106 (2020).

21. Fane, M. E. et al. Stromal changes in the aged lung induce an emergence from melanoma dormancy. Nature 606, 396–405 (2022).

22. Johnson, M. et al. Advanced Age in Humans and Mouse Models of Glioblastoma Show Decreased Survival from Extratumoral Influence. Clinical Cancer Research 29, 4973–4989 (2023).

23. Turrell, F. K. et al. Age-associated microenvironmental changes highlight the role of PDGF-C in ER+ breast cancer metastatic relapse. Nat Cancer 4, 468–484 (2023).

24. Bianchi-Frias, D. et al. The Aged Microenvironment Influences the Tumorigenic Potential of Malignant Prostate Epithelial Cells. Mol Cancer Res 17, 321–331 (2019).

25. Alicea, G. M. et al. Changes in Aged Fibroblast Lipid Metabolism Induce Age-Dependent Melanoma Cell Resistance to Targeted Therapy via the Fatty Acid Transporter FATP2. Cancer Discov 10, 1282–1295 (2020).

26. Gupta, P. et al. The Aging Microenvironment is a Determinant of Immune Exclusion and Metastatic Fate in Pancreatic Cancer. Cancer Res 10.1158/0008-5472.CAN-25-1904 (2025) doi:10.1158/0008-5472.CAN-25-1904.

27. Zabransky, D. J. et al. Fibroblasts in the Aged Pancreas Drive Pancreatic Cancer Progression. Cancer Res 84, 1221–1236 (2024).

28. Zhang, Z. et al. A panoramic view of cell population dynamics in mammalian aging. Science 387, eadn3949 (2025).

29. Oni, T. E. et al. SOAT1 promotes mevalonate pathway dependency in pancreatic cancer. J Exp Med 217, (2020).

30. Boj, S. F. et al. Organoid models of human and mouse ductal pancreatic cancer. Cell 160, 324–338 (2015).

31. Hingorani, S. R. et al. Trp53R172H and KrasG12D cooperate to promote chromosomal instability and widely metastatic pancreatic ductal adenocarcinoma in mice. Cancer Cell 7, 469–83 (2005).

32. Mucciolo, G. et al. EGFR-activated myofibroblasts promote metastasis of pancreatic cancer. Cancer Cell 10.1016/j.ccell.2023.12.002 (2023) doi:10.1016/j.ccell.2023.12.002.

33. Tsang, E. S. et al. Delving into Early-onset Pancreatic Ductal Adenocarcinoma: How Does Age Fit In? Clin Cancer Res 27, 246–254 (2021).

34. Elyada, E. et al. Cross-species single-cell analysis of pancreatic ductal adenocarcinoma reveals antigen-presenting cancer-associated fibroblasts. Cancer Discov 10.1158/2159-8290.CD-19-0094 (2019) doi:10.1158/2159-8290.CD-19-0094.

35. Lloyd, E. G. et al. SMAD4 and KRAS Status Shape Cancer Cell-Stromal Crosstalk and Therapeutic Response in Pancreatic Cancer. Cancer Research 10.1158/0008-5472.CAN-24-2330 (2025) doi:10.1158/0008-5472.CAN-24-2330.

36. Biffi, G. et al. IL1-Induced JAK/STAT Signaling Is Antagonized by TGFbeta to Shape CAF Heterogeneity in Pancreatic Ductal Adenocarcinoma. Cancer Discov 9, 282–301 (2019).

37. Bianchi, A. et al. Cell-Autonomous Cxcl1 Sustains Tolerogenic Circuitries and Stromal Inflammation via Neutrophil-Derived TNF in Pancreatic Cancer. Cancer Discovery 13, 1428–1453 (2023).

38. Cumming, J. et al. Dissecting FAP+ Cell Diversity in Pancreatic Cancer Uncovers an Interferon-Response Subtype of Cancer-Associated Fibroblasts with Tumor-Restraining Properties. Cancer Res 85, 2388–2411 (2025).

39. Jassim, A. et al. Gene context drift identifies drug targets to mitigate cancer treatment resistance. Cancer Cell S1535–6108(25)00255–7 (2025) doi:10.1016/j.ccell.2025.06.005.

40. Chijimatsu, R. et al. Establishment of a reference single-cell RNA sequencing dataset for human pancreatic adenocarcinoma. iScience 25, 104659 (2022).

41. Tyner, J. W. et al. Understanding Drug Sensitivity and Tackling Resistance in Cancer. Cancer Res 82, 1448–1460 (2022).

42. Cannon, M. et al. DGIdb 5.0: rebuilding the drug-gene interaction database for precision medicine and drug discovery platforms. Nucleic Acids Res 52, D1227–D1235 (2024).

43. di Micco, P. et al. canSAR: update to the cancer translational research and drug discovery knowledgebase. Nucleic Acids Res 51, D1212–D1219 (2023).

44. Mukherjee, S. et al. Toll-like receptor-guided therapeutic intervention of human cancers: molecular and immunological perspectives. Front Immunol 14, 1244345 (2023).

45. Somani, V. K. et al. IRAK4 Signaling Drives Resistance to Checkpoint Immunotherapy in Pancreatic Ductal Adenocarcinoma. Gastroenterology 162, 2047–2062 (2022).

46. Zhang, D. et al. Constitutive IRAK4 Activation Underlies Poor Prognosis and Chemoresistance in Pancreatic Ductal Adenocarcinoma. Clin Cancer Res 23, 1748–1759 (2017).

47. Zhang, D. et al. Tumor-Stroma IL1β-IRAK4 Feedforward Circuitry Drives Tumor Fibrosis, Chemoresistance, and Poor Prognosis in Pancreatic Cancer. Cancer Res 78, 1700–1712 (2018).

48. Matsui, H. et al. Survival Analysis of 4 Different Age Groups of Pancreatic Ductal Adenocarcinoma After Radical Resection From Retrospective Multi-Center Analysis (YPB-003). Cancer Med 14, e70647 (2025).

49. Høj, K. et al. Age-Related Decline in Pancreas Regeneration Is Associated With an Increased Proinflammatory Response to Injury. Gastro Hep Adv 3, 973–985 (2024).

50. Dube, C. T. et al. Age-Related Alterations in Macrophage Distribution and Function Are Associated With Delayed Cutaneous Wound Healing. Front Immunol 13, 943159 (2022).

51. Moss, C. E. et al. Aging-related defects in macrophage function are driven by MYC and USF1 transcriptional programs. Cell Rep 43, 114073 (2024).

52. Ma, R.-Y., Black, A. & Qian, B.-Z. Macrophage diversity in cancer revisited in the era of single-cell omics. Trends Immunol 43, 546–563 (2022).

53. Shah, Y. et al. Pan-cancer analysis reveals molecular patterns associated with age. Cell Rep 37, 110100 (2021).

54. Di Micco, R., Krizhanovsky, V., Baker, D. & d’Adda di Fagagna, F. Cellular senescence in ageing: from mechanisms to therapeutic opportunities. Nat Rev Mol Cell Biol 22, 75–95 (2021).

55. Belle, J. I. et al. Senescence Defines a Distinct Subset of Myofibroblasts That Orchestrates Immunosuppression in Pancreatic Cancer. Cancer Discov 14, 1324–1355 (2024).

56. Ye, J. et al. Senescent CAFs Mediate Immunosuppression and Drive Breast Cancer Progression. Cancer Discov 14, 1302–1323 (2024).

57. Assouline, B. et al. Senescent cancer-associated fibroblasts in pancreatic adenocarcinoma restrict CD8+ T cell activation and limit responsiveness to immunotherapy in mice. Nat Commun 15, 6162 (2024).

58. Zhuang, X. et al. Ageing limits stemness and tumorigenesis by reprogramming iron homeostasis. Nature 637, 184–194 (2025).

59. Gyawali, B. & Booth, C. M. Treatment of metastatic pancreatic cancer: 25 years of innovation with little progress for patients. Lancet Oncol 25, 167–170 (2024).

60. Thota, R., Maitra, A. & Berlin, J. D. Preclinical Rationale for the Phase III Trials in Metastatic Pancreatic Cancer: Is Wishful Thinking Clouding Successful Drug Development for Pancreatic Cancer? Pancreas 46, 143–150 (2017).

61. Dasgupta, A., et al. Anticachectic regulator analysis reveals Perp-dependent antitumorigenic properties of 3-methyladenine in pancreatic cancer. JCI Insight 7, e153842 (2022).

62. Ogobuiro, I., et al. Multiomic Characterization Reveals a Distinct Molecular Landscape in Young-Onset Pancreatic Cancer. JCO Precis Oncol 7, e2300152 (2023).

63. Ben-Aharon, I. et al. Genomic Landscape of Pancreatic Adenocarcinoma in Younger versus Older Patients: Does Age Matter? Clin Cancer Res 25, 2185–2193 (2019).

64. Li, Q., et al. IRAK4 mediates colitis-induced tumorigenesis and chemoresistance in colorectal cancer. JCI Insight 4, (2019).

65. Von Roemeling, C. A. et al. Oral IRAK-4 Inhibitor CA-4948 Is Blood-Brain Barrier Penetrant and Has Single-Agent Activity against CNS Lymphoma and Melanoma Brain Metastases. Clinical Cancer Research 29, 1751–1762 (2023).

66. Guidetti, F. et al. Targeting IRAK4 with Emavusertib in Lymphoma Models with Secondary Resistance to PI3K and BTK Inhibitors. J Clin Med 12, 399 (2023).

67. Cancer Genome Atlas Research Network. Electronic address: & Cancer Genome Atlas Research Network. Integrated Genomic Characterization of Pancreatic Ductal Adenocarcinoma. Cancer Cell 32, 185-203.e13 (2017).

68. Mahat, D. B. et al. Mutant p53 exploits enhancers to elevate immunosuppressive chemokine expression and impair immune checkpoint inhibitors in pancreatic cancer. Immunity 58, 1688–1705.e9 (2025).

69. Vennin, C. et al. CAF hierarchy driven by pancreatic cancer cell p53-status creates a pro-metastatic and chemoresistant environment via perlecan. Nat Commun 10, 3637 (2019).

70. Maddalena, M. et al. TP53 missense mutations in PDAC are associated with enhanced fibrosis and an immunosuppressive microenvironment. Proc Natl Acad Sci U S A 118, e2025631118 (2021).

71. Jihad, M. et al. Protocol for the characterization of the pancreatic tumor microenvironment using organoid-derived mouse models and single-nuclei RNA sequencing. STAR Protoc 5, 103203 (2024).

72. Gummadi, V. R. et al. Discovery of CA-4948, an Orally Bioavailable IRAK4 Inhibitor for Treatment of Hematologic Malignancies. ACS Med Chem Lett 11, 2374–2381 (2020).

73. Ohlund, D. et al. Distinct populations of inflammatory fibroblasts and myofibroblasts in pancreatic cancer. J Exp Med 214, 579–596 (2017).

74. Humpton, T. J. et al. Oncogenic KRAS Induces NIX-Mediated Mitophagy to Promote Pancreatic Cancer. Cancer Discovery 9, 1268–1287 (2019).

75. Zheng, G. X. Y. et al. Massively parallel digital transcriptional profiling of single cells. Nat Commun 8, 14049 (2017).

76. Fleming, S. J. et al. Unsupervised removal of systematic background noise from droplet-based single-cell experiments using CellBender. Nat Methods 20, 1323–1335 (2023).

77. Wolf, F. A., Angerer, P. & Theis, F. J. SCANPY: large-scale single-cell gene expression data analysis. Genome Biology 19, 15 (2018).

78. Bernstein, N. J. et al. Solo: Doublet Identification in Single-Cell RNA-Seq via Semi-Supervised Deep Learning. Cell Syst 11, 95–101.e5 (2020).

79. Lange, M. et al. CellRank for directed single-cell fate mapping. Nat Methods 19, 159–170 (2022).

80. Gulati, G. S. et al. Single-cell transcriptional diversity is a hallmark of developmental potential. Science 367, 405–411 (2020).

81. Finak, G. et al. MAST: a flexible statistical framework for assessing transcriptional changes and characterizing heterogeneity in single-cell RNA sequencing data. Genome Biology 16, 278 (2015).

82. Subramanian, A. et al. Gene set enrichment analysis: A knowledge-based approach for interpreting genome-wide expression profiles. Proceedings of the National Academy of Sciences 102, 15545–15550 (2005).

83. Yu, G. Thirteen years of clusterProfiler. The Innovation 5, 100722 (2024).

84. Liberzon, A. et al. The Molecular Signatures Database Hallmark Gene Set Collection. Cell Systems 1, 417–425 (2015).

85. Peng, J. et al. Single-cell RNA-seq highlights intra-tumoral heterogeneity and malignant progression in pancreatic ductal adenocarcinoma. Cell Res 29, 725–738 (2019).

86. Lin, W. et al. Single-cell transcriptome analysis of tumor and stromal compartments of pancreatic ductal adenocarcinoma primary tumors and metastatic lesions. Genome Med 12, 80 (2020).

87. Halbrook, C. J. et al. Differential integrated stress response and asparagine production drive symbiosis and therapy resistance of pancreatic adenocarcinoma cells. Nat Cancer 3, 1386–1403 (2022).

88. Korsunsky, I. et al. Fast, sensitive and accurate integration of single-cell data with Harmony. Nat Methods 16, 1289–1296 (2019).

89. Traag, V. A., Waltman, L. & van Eck, N. J. From Louvain to Leiden: guaranteeing well-connected communities. Sci Rep 9, 5233 (2019).

90. Grover, A. & Leskovec, J. node2vec: Scalable Feature Learning for Networks. KDD 2016, 855–864 (2016).

91. Mikolov, T., Chen, K., Corrado, G. & Dean, J. Efficient Estimation of Word Representations in Vector Space. Preprint at 10.48550/arXiv.1301.3781 (2013).

92. Hagberg, A. A., Schult, D. A. & Swart, P. J. Exploring Network Structure, Dynamics, and Function using NetworkX. scipy 10.25080/TCWV9851 (2008) doi:10.25080/TCWV9851.

93. Kamada, T. & Kawai, S. An algorithm for drawing general undirected graphs. Information Processing Letters 31, 7–15 (1989).

94. Hao, Y. et al. Dictionary learning for integrative, multimodal and scalable single-cell analysis. Nat Biotechnol 42, 293–304 (2024).

95. Patro, R., Duggal, G., Love, M. I., Irizarry, R. A. & Kingsford, C. Salmon provides fast and bias-aware quantification of transcript expression. Nat Methods 14, 417–419 (2017).

96. Soneson, C., Love, M. I. & Robinson, M. D. Differential analyses for RNA-seq: transcript-level estimates improve gene-level inferences. F1000Res 4, 1521 (2015).

97. Love, M. I., Huber, W. & Anders, S. Moderated estimation of fold change and dispersion for RNA-seq data with DESeq2. Genome Biology 15, 550 (2014).

98. Zhu, A., Ibrahim, J. G. & Love, M. I. Heavy-tailed prior distributions for sequence count data: removing the noise and preserving large differences. Bioinformatics 35, 2084 (2019).

99. Mauri, M., Elli, T., Caviglia, G., Uboldi, G. & Azzi, M. RAWGraphs: A Visualisation Platform to Create Open Outputs. in Proceedings of the 12th Biannual Conference on Italian SIGCHI Chapter 1–5 (Association for Computing Machinery, New York, NY, USA, 2017). doi:10.1145/3125571.3125585.

100. Büttner, M., Ostner, J., Müller, C. L., Theis, F. J. & Schubert, B. scCODA is a Bayesian model for compositional single-cell data analysis. Nat Commun 12, 6876 (2021).

101. Dominguez, C. X. et al. Single-Cell RNA Sequencing Reveals Stromal Evolution into LRRC15(+) Myofibroblasts as a Determinant of Patient Response to Cancer Immunotherapy. Cancer Discov 10, 232–253 (2020).

102. McAndrews, K. M. et al. Identification of Functional Heterogeneity of Carcinoma-Associated Fibroblasts with Distinct IL6-Mediated Therapy Resistance in Pancreatic Cancer. Cancer Discov 12, 1580–1597 (2022).

