## Supplemental Figures for "Cross-species graph-embedding unmasks the ageing microenvironment as a key determinant of pancreatic cancer malignant cell biology and therapy response"

### EXTENDED DATA FIGURES AND FIGURE LEGENDS

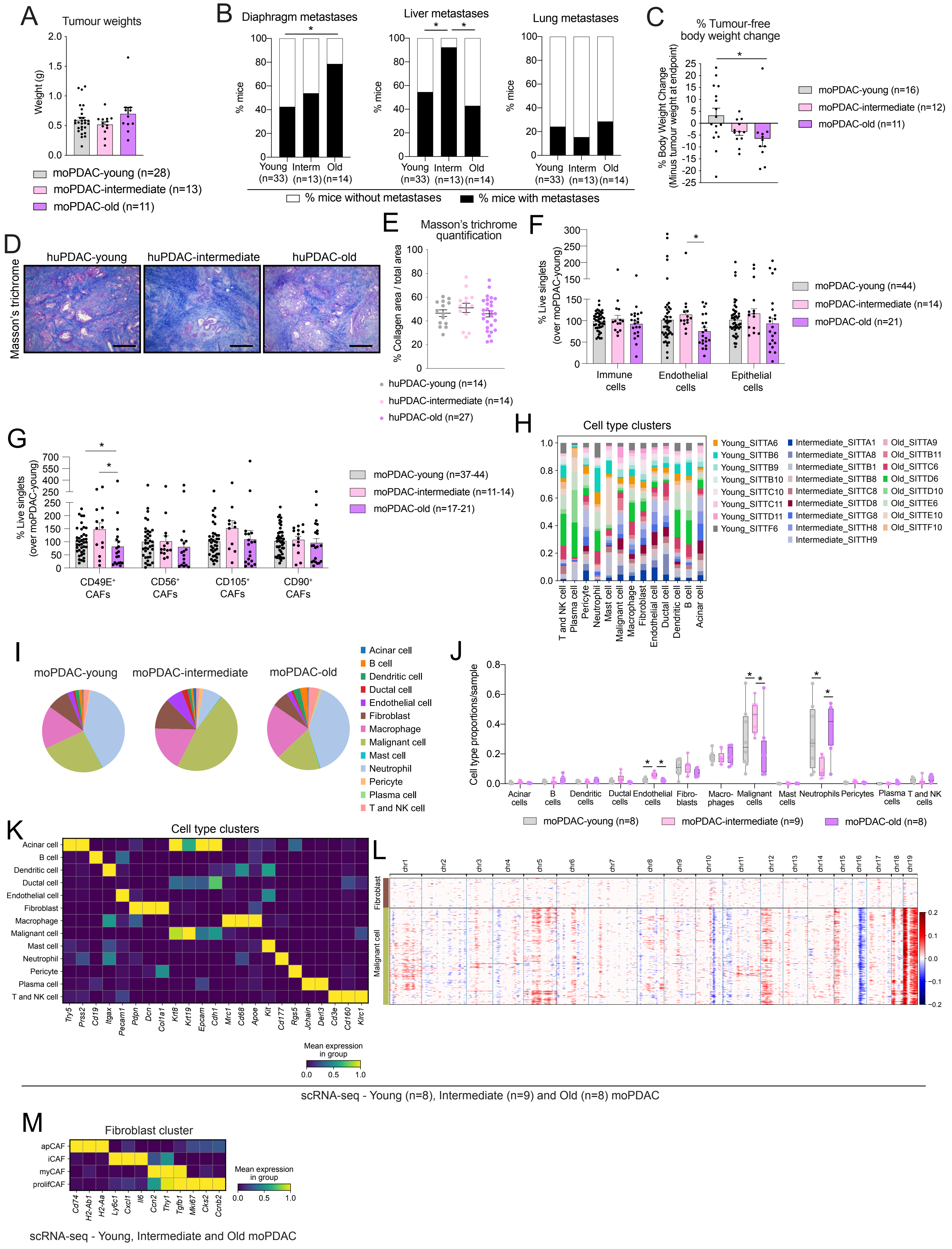


Extended Data Fig. 1

**Extended Data Figure 1. Ageing promotes an inflammatory CAF phenotype in PDAC mouse models.** **(A)** Tumour weights measured at experimental endpoint of young (~2-3-month-old, moPDAC-young), intermediate (~12-13-month-old, moPDAC-intermediate) and old (~18-20-month-old, moPDAC-old) mice. Results show mean ± SEM. No statistical difference was found, as calculated by Kruskal-Wallis test. **(B)** moPDAC-young, moPDAC-intermediate and moPDAC-old mice with or without metastases in the diaphragm, liver, and lungs taken at comparable endpoints. *, $P < 0.05$, Fisher's test. **(C)** Body weight change in moPDAC-young, moPDAC-intermediate and moPDAC-old at endpoint compared to body weight at surgery. Results show mean ± SEM. *, $P\ adj < 0.05$, Kruskal-Wallis test. Tumour-free body weight was calculated by removing the tumour weight from the total mouse weight at endpoint. **(D)** Representative Masson's trichrome stains in young (≤ 55-year-old; hereon, huPDAC-young), intermediate (56–69-year-old; huPDAC-intermediate) and old (≥ 70-year-old; huPDAC-old) human (hu)PDAC tumours. Scale bars, 200 μm. **(E)** Quantification of Masson's trichrome stain in huPDAC-young, huPDAC-intermediate and huPDAC-old. Results show mean ± SEM. No statistical difference was found, as calculated by Kruskal-Wallis test. **(F)** Flow cytometric analysis of immune cells ($CD45^+CD31^-EpCAM^-$), endothelial cells ($CD31^+CD45^-EpCAM^-$) and epithelial cells ($CD45^-CD31^-EpCAM^+$) from live singlets in moPDAC-young, moPDAC-intermediate and moPDAC-old. Results show mean ± SEM. *, $P\ adj < 0.05$, Kruskal-Wallis test. **(G)** Flow cytometric analyses of $CD49E^+$, $CD56^+$, $CD105^+$ and $CD90^+$ cancer-associated fibroblast (CAF) populations from live singlets in moPDAC-young, moPDAC-intermediate and moPDAC-old. Results show mean ± SEM. *, $P\ adj < 0.05$, Kruskal-Wallis test. **(H)** Tumour sample contribution to different cell types in moPDAC-young (n=8), moPDAC-intermediate (n=9) and moPDAC-old (n=8), represented as bar plot showing proportions of different colour-coded tumour samples in each cell cluster of the combined moPDAC single cell RNA-sequencing (scRNA-seq) datasets. **(I)** Cell type contribution in moPDAC-young, moPDAC-intermediate and moPDAC-old represented as pie charts showing proportions of the different cell clusters in each condition. **(J)** Boxplots illustrating the proportion of cell types across samples stratified by age group (moPDAC-young, moPDAC-intermediate and moPDAC-old). Statistically credible changes in cell types across age groups are denoted with *, as tested with sCODA, with FDR < 0.05. **(K)** Heatmap of scaled expression of cell type-specific markers in each cell cluster of the combined scRNA-seq. Data are scaled such that the cluster with the lowest average expression = 0 and the highest = 1 for each gene. **(L)** Heatmap showing large-scale copy number variation (CNV) profile of the fibroblast and epithelial malignant cell clusters identified by InferCNV analysis of the combined moPDAC scRNA-seq datasets. The colour coding represents the CNV level based on a sliding window of 250 gene expression. Amplifications are shown in red, and deletions are shown in blue. The fibroblast cluster was used as reference cell cluster. **(M)** Heatmap of scaled expression of CAF-specific markers in each CAF cluster of the combined moPDAC scRNA-seq datasets. Data are scaled such that the cluster with the lowest average expression = 0 and the highest = 1 for each gene.

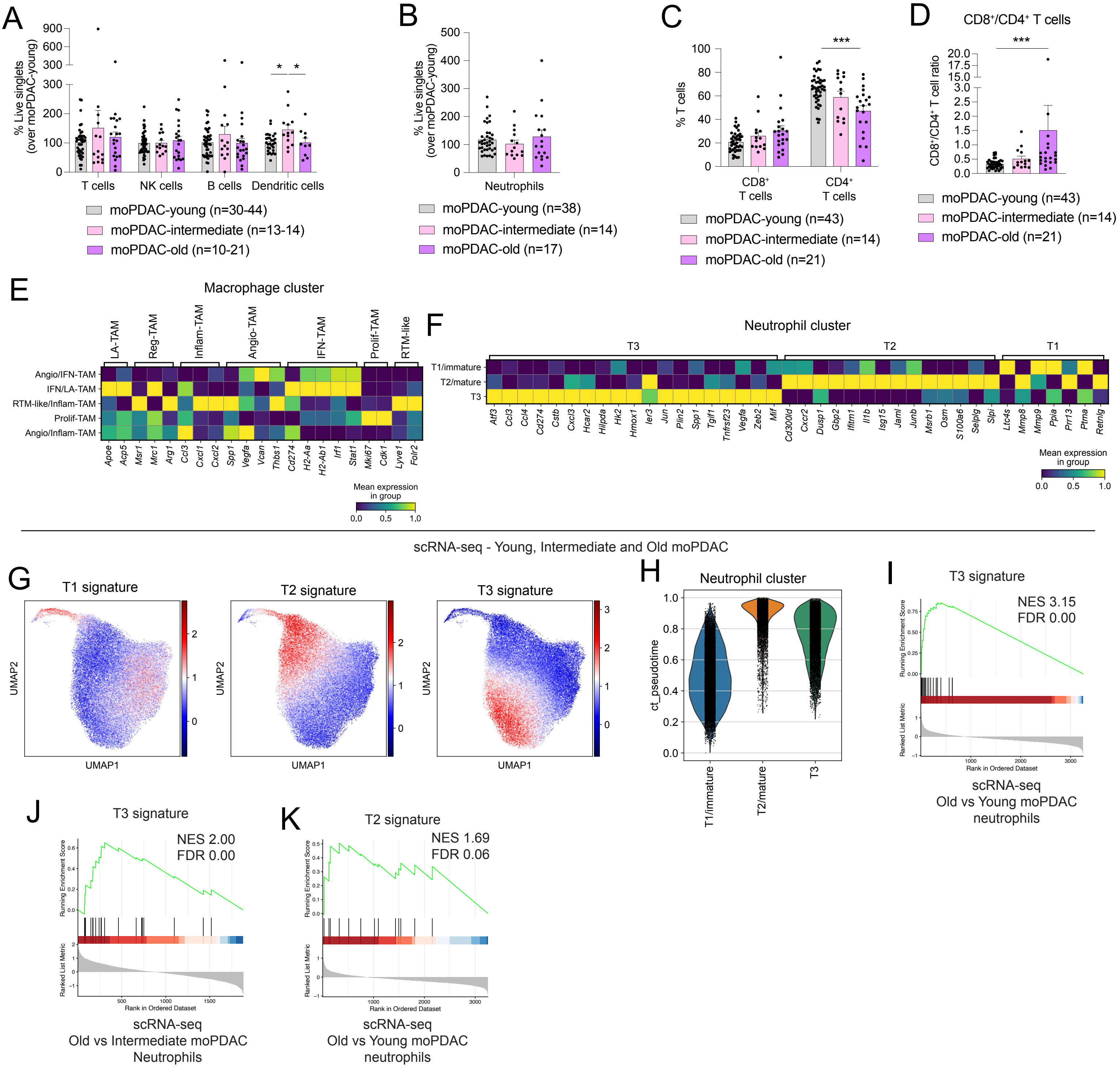

A
% Live singlets (over moPDAC-young)
T cells
NK cells
B cells
Dendritic cells
moPDAC-young (n=30-44)
moPDAC-intermediate (n=13-14)
moPDAC-old (n=10-21)
B
% Live singlets (over moPDAC-young)
Neutrophils
moPDAC-young (n=38)
moPDAC-intermediate (n=14)
moPDAC-old (n=17)
C
% T cells
CD8+ T cells
CD4+ T cells
moPDAC-young (n=43)
moPDAC-intermediate (n=14)
moPDAC-old (n=21)
D
CD8+/CD4+ T cells
CD8+/CD4+ T cell ratio
moPDAC-young (n=43)
moPDAC-intermediate (n=14)
moPDAC-old (n=21)
E
Macrophage cluster
LA-TAM
Reg-TAM
Inflam-TAM
Angio-TAM
IFN-TAM
Prolif-TAM
RTM-like
Angio/IFN-TAM
IFN/LA-TAM
RTM-like/Inflam-TAM
Prolif-TAM
Angio/Inflam-TAM
Apoe Acp5 Msr1 Mrc1 Arg1 Ccl3 Cxcl1 Cxcl2 Spp1 Vegfa Vcan Thbs1 Cd274 H2-Aa H2-Ab1 Irf1 Stat1 Mki67 Cdk1 Lyve1 Folr2
Mean expression in group
F
Neutrophil cluster
T3
T2
T1
T1/immature
T2/mature
T3
Atf3 Ccl3 Ccl4 Cd274 Cstb Cxcl3 Hcar2 Hilpda Hk2 Hmox1 Ier3 Jun Plin2 Spp1 Tgif1 Tnfrsf23 Vegfa Zeb2 Mif Cd300ld Cxcr2 Dusp1 Gbp2 Ifitm1 Il1b Isg15 Jaml Junb Msrb1 Osm S100a6 Selplg Slpi Ltc4s Mmp8 Mmp9 Ppia Prr13 Ptma Retnlg
Mean expression in group
scRNA-seq - Young, Intermediate and Old moPDAC
G
T1 signature
T2 signature
T3 signature
UMAP1
UMAP2
H
Neutrophil cluster
ct_pseudotime
T1/immature
T2/mature
T3
I
T3 signature
NES 3.15
FDR 0.00
Running Enrichment Score
Ranked List Metric
Rank in Ordered Dataset
scRNA-seq
Old vs Young moPDAC
neutrophils
J
T3 signature
NES 2.00
FDR 0.00
scRNA-seq
Old vs Intermediate moPDAC
Neutrophils
K
T2 signature
NES 1.69
FDR 0.06
scRNA-seq
Old vs Young moPDAC
neutrophils

**Extended Data Figure 2. Ageing shapes immune cell composition in PDAC mouse models. (A)** Flow cytometric analysis of T cells ($CD45^+CD3\varepsilon^+TCR\beta^+$), Natural Killer (NK) cells ($CD45^+NK1.1^+$), B cells ($CD45^+CD19^+$) and dendritic cells ($CD45^+$ $CD11b^+Gr1^-CD11c^+$) from live singlets in moPDAC-young, moPDAC-intermediate and moPDAC-old. Results show mean ± SEM. *, *P adj* < 0.05, Kruskal-Wallis test. **(B)** Flow cytometric analysis of neutrophils ($CD45^+CD11b^+Gr1^+$) from live singlets in moPDAC-young, moPDAC-intermediate and moPDAC-old. Results show mean ± SEM. No statistical difference was found, as calculated by Kruskal-Wallis test. **(C)** Flow cytometric analysis of $CD4^+$ and $CD8^+$ T cells ($CD45^+CD3\varepsilon^+TCR\beta^+$) from the parental gate in moPDAC-young, moPDAC-intermediate and moPDAC-old. Results show mean ± SEM. ***, *P adj* < 0.001, Kruskal-Wallis test. **(D)** $CD8^+/CD4^+$ T cell ratio, as assessed by flow cytometry from the parental gate in moPDAC-young, moPDAC-intermediate and moPDAC-old. Results show mean ± SEM. ***, *P adj* < 0.001, Kruskal-Wallis test. **(E)** Heatmap of scaled expression of macrophage subset markers in distinct macrophage sub-clusters of the combined scRNA-seq datasets. Data are scaled such that the cluster with the lowest average expression = 0 and the highest = 1 for each gene. **(F)** Heatmap of scaled expression of neutrophil subset markers in distinct neutrophil sub-clusters of the combined scRNA-seq datasets. Data are scaled such that the cluster with the lowest average expression = 0 and the highest = 1 for each gene. **(G)** Uniform manifold approximation and projection (UMAP) plots highlighting the enrichment of T1, T2, T3 neutrophil gene signatures in the neutrophil cluster of the combined scRNA-seq datasets. The signatures are from Ng et al. **(H)** Violin plots showing the distribution of ct_pseudotime scores, computed using the CellRank CytoTRACE kernel (inverse of the CytoTRACE score), across cells within each neutrophil subset. The T1/immature subset exhibits the lowest ct_pseudotime values, indicative of a less differentiated phenotype. **(I-J)** GSEA of T3 gene signature in neutrophils from moPDAC-old compared to neutrophils from moPDAC-young **(I)** and in neutrophils from moPDAC-old compared to neutrophils from moPDAC-intermediate **(J)**, as assessed by MAST analysis of the scRNA-seq dataset. The signature was significantly altered. **(K)** GSEA of T2 gene signature in neutrophils from moPDAC-old compared to neutrophils from moPDAC-young, as assessed by MAST analysis of the scRNA-seq dataset. The signature was significantly altered.

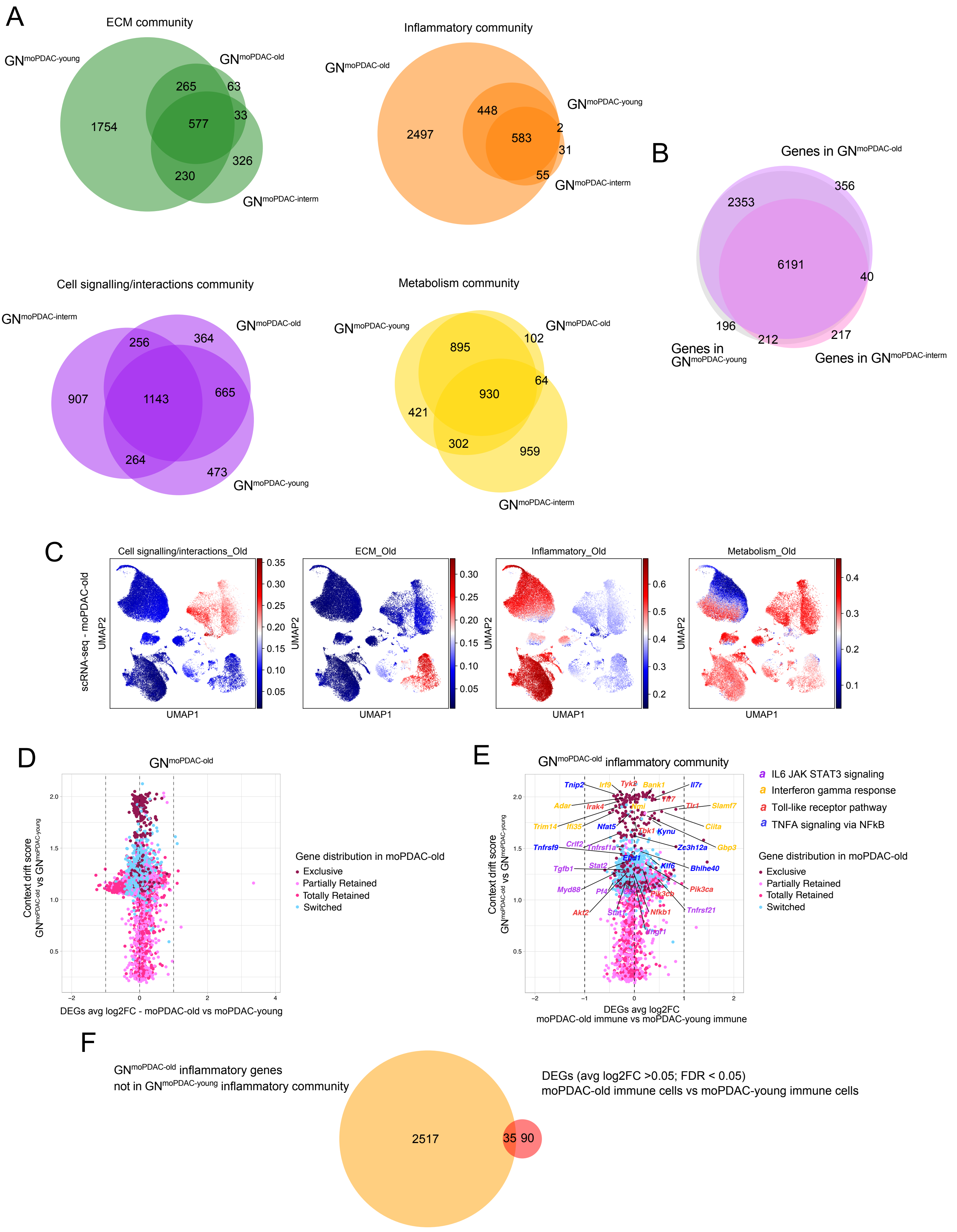


Extended Data Fig. 3

**Extended Data Figure 3. Graph-embedding reveals age-dependent acquisition of inflammatory functions in PDAC mouse models. (A)** Venn diagrams showing the overlap between each gene community in *RE*sistance through *CO*ntext *DR*ift (RECODR) graph networks (GNs) of moPDAC-young (hereon, GN$^{\text{moPDAC-young}}$), moPDAC-intermediate (hereon, GN$^{\text{moPDAC-interm}}$) and moPDAC-old (hereon, GN$^{\text{moPDAC-old}}$). **(B)** Venn diagram showing the overlap between genes in GN$^{\text{moPDAC-young}}$, GN$^{\text{moPDAC-interm}}$ and GN$^{\text{moPDAC-old}}$. **(C)** UMAP plots of all cell type clusters from moPDAC scRNA-seq highlighting the enrichment of GN$^{\text{moPDAC-old}}$ communities. **(D)** X-Y scatter plot of all GN$^{\text{moPDAC-old}}$ genes with context drift score (CDS) that are DEGs (FDR < 0.05) for moPDAC-old compared to moPDAC-young (n=8,801), as assessed by MAST analysis. Dotted lines mark the DEG average Log2FC threshold (avg Log2FC > 1 or < -1). Dots (representing genes) are coloured based on their gene distribution on GN$^{\text{moPDAC-old}}$ as 'exclusive', 'switched', 'partially retained' or 'totally retained'. **(E)** X-Y scatter plot of GN$^{\text{moPDAC-old}}$ inflammatory genes with CDS that are DEGs (FDR < 0.05) for moPDAC-old immune cells (comprising macrophages, neutrophils and dendritic cells) compared to moPDAC-young immune cells (n=2,802), as assessed by MAST analysis. Dotted lines mark the avg DEG Log2FC threshold (avg Log2FC > 1 or < -1). Dots (representing genes) are colour coded based on their gene distribution on moPDAC-old as 'exclusive', 'switched', 'partially retained' or 'totally retained'. Selected gene names are coloured based on their presence in pathways enriched in the GN$^{\text{moPDAC-old}}$ inflammatory community. **(F)** Venn diagram showing the overlap between GN$^{\text{moPDAC-old}}$ inflammatory genes not present in the GN$^{\text{moPDAC-young}}$ inflammatory community and DEGs with avg Log2FC > 0.05, FDR < 0.05 of moPDAC-old immune cells (comprising macrophages, neutrophils and dendritic cells) compared to moPDAC-young immune cells, as assessed by MAST analysis.

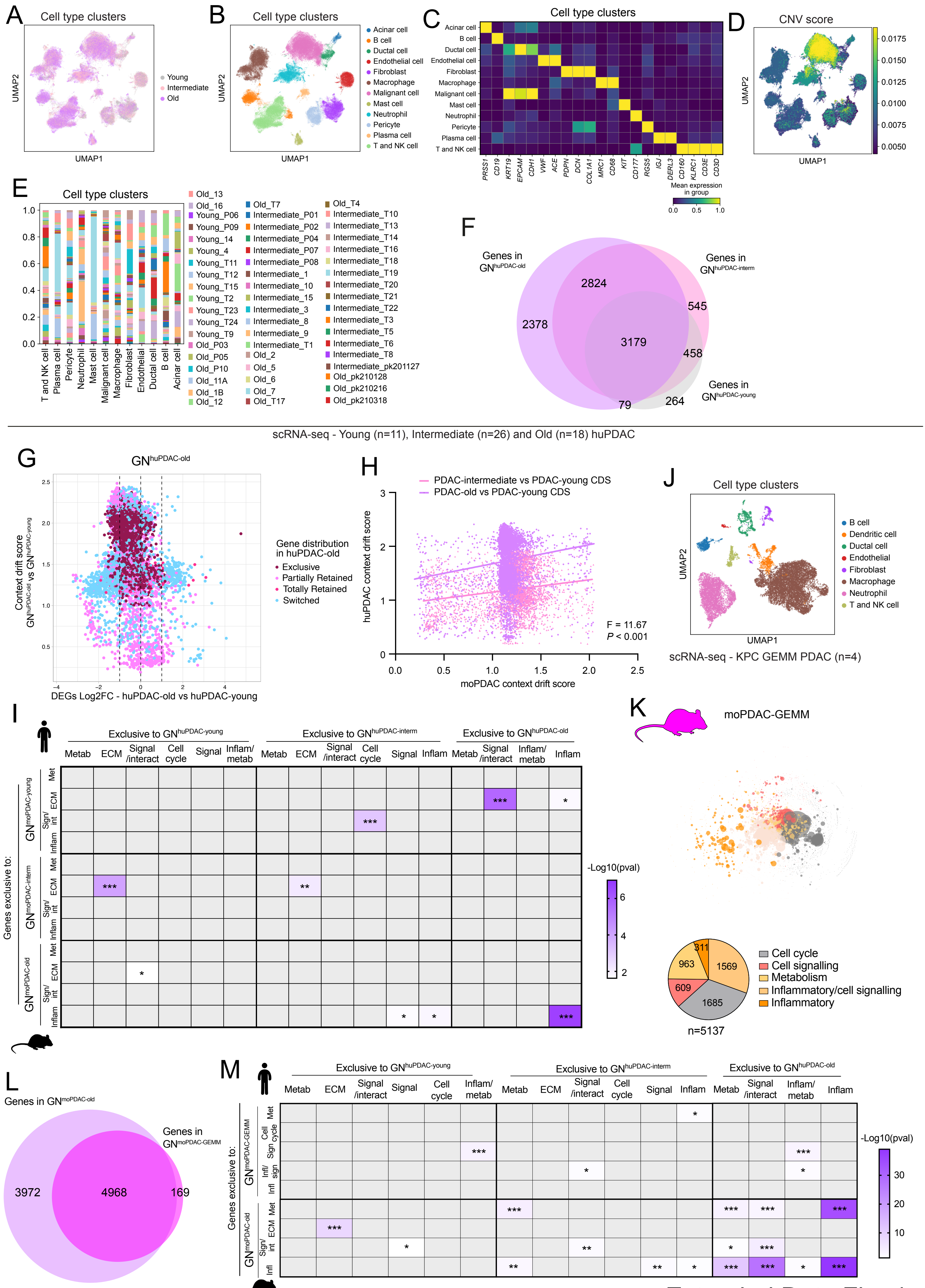


Extended Data Fig. 4

**Extended Data Figure 4. Old PDAC mouse models capture age-specific changes of human PDAC. (A-B)** UMAP plots of all cells from huPDAC-young (n=11), huPDAC-intermediate (n=26) and huPDAC-old (n=18) analysed by scRNA-seq. Different ages **(A)** or cell types **(B)** are colour coded. scRNA-seq data are from Chijimatsu et al. **(C)** Heatmap of scaled expression of cell type-specific markers in each cell cluster of the combined huPDAC scRNA-seq datasets. Data are scaled such that the cluster with the lowest average expression = 0 and the highest = 1 for each gene. **(D)** UMAP showing the inferred CNV score of all cell clusters identified by scRNA-seq of huPDAC. **(E)** Tumour sample contribution to different cell types in the combined huPDAC scRNA-seq datasets represented as bar plots showing proportions of the different colour coded tumour samples in each cell cluster. **(F)** Venn diagram showing the overlap between genes in GNs of huPDAC-young (hereon, GN$^{huPDAC\text{-}young}$), huPDAC-intermediate (hereon, GN$^{huPDAC\text{-}interm}$) and huPDAC-old (hereon, GN$^{huPDAC\text{-}old}$). **(G)** X-Y scatter plot of all GN$^{huPDAC\text{-}old}$ genes with CDS that are DEGs (FDR < 0.05) for huPDAC-old compared to huPDAC-young (n=7,719), as assessed by MAST analysis of the scRNA-seq data. Dotted lines mark the DEG Log2FC threshold (Log2FC > 1 or < -1). Dots (representing genes) are colour coded based on their gene distribution on GN$^{huPDAC\text{-}old}$ as 'exclusive', 'switched', 'partially retained' or 'totally retained'. **(H)** Linear regression between CDS values of GN$^{moPDAC}$ and GN$^{huPDAC}$. Cross-species comparisons across common genes between PDAC-old and PDAC-young CDS values (purple; n=5,869; slope = 0.4 ± 0.03; R squared value = 0.02) and across common genes between PDAC-intermediate and PDAC-young CDS values (pink; n=3,783; slope = 0.2 ± 0.02; R squared value = 0.03) are shown. The difference between the two slopes was significant, as assessed by F-test (F = 11.67; *P* = 0.0006). **(I)** Heatmap showing the overlap of 'exclusive' genes in each community on GN$^{moPDAC\text{-}young}$, GN$^{moPDAC\text{-}interm}$ and GN$^{moPDAC\text{-}old}$ with 'exclusive' genes in each community on GN$^{huPDAC\text{-}young}$, GN$^{huPDAC\text{-}interm}$ and GN$^{huPDAC\text{-}old}$. Communities without any 'exclusive' genes were not included in this analysis. Grey spaces indicate not significant overlaps. Significance was defined by hypergeometric test with representation factor > 1; * *P* < 0.05, ** *P* < 0.01, *** *P* < 0.001; denominator = 17,094 orthologs. **(J)** UMAP plot of all cells from KPC (*Kras*$^{LSL\text{-}G12D/+}$; *Trp53*$^{LSL\text{-}R172H/+}$; *Pdx1-Cre*) genetically engineered mouse model (GEMM)-derived PDAC tumours (n=4), analysed by scRNA-seq. Different cell types are colour coded. scRNA-seq data are from Elyada et al. **(K)** RECODR GN (top) and pie chart (bottom) of gene communities generated from scRNA-seq datasets of KPC GEMM-derived PDAC tumours (n=4; hereon, GN$^{moPDAC\text{-}GEMM}$; n=5,137 genes; n=15,075 cells). **(L)** Venn diagram showing the overlap between genes in GN$^{moPDAC\text{-}GEMM}$ and GN$^{moPDAC\text{-}old}$. **(M)**

Heatmap showing the overlap of ‘exclusive’ genes in each community on $GN^{moPDAC\text{-}young}$ and $GN^{moPDAC\text{-}GEMM}$ with ‘exclusive’ genes in each community on $GN^{huPDAC\text{-}young}$, $GN^{huPDAC\text{-}interm}$ and $GN^{huPDAC\text{-}old}$. Communities without any ‘exclusive’ genes were not included in this analysis. Grey spaces indicate not significant overlaps. Significance was defined by hypergeometric test with representation factor > 1; * $P < 0.05$, ** $P < 0.01$, *** $P < 0.001$; denominator = 17,094 orthologs.

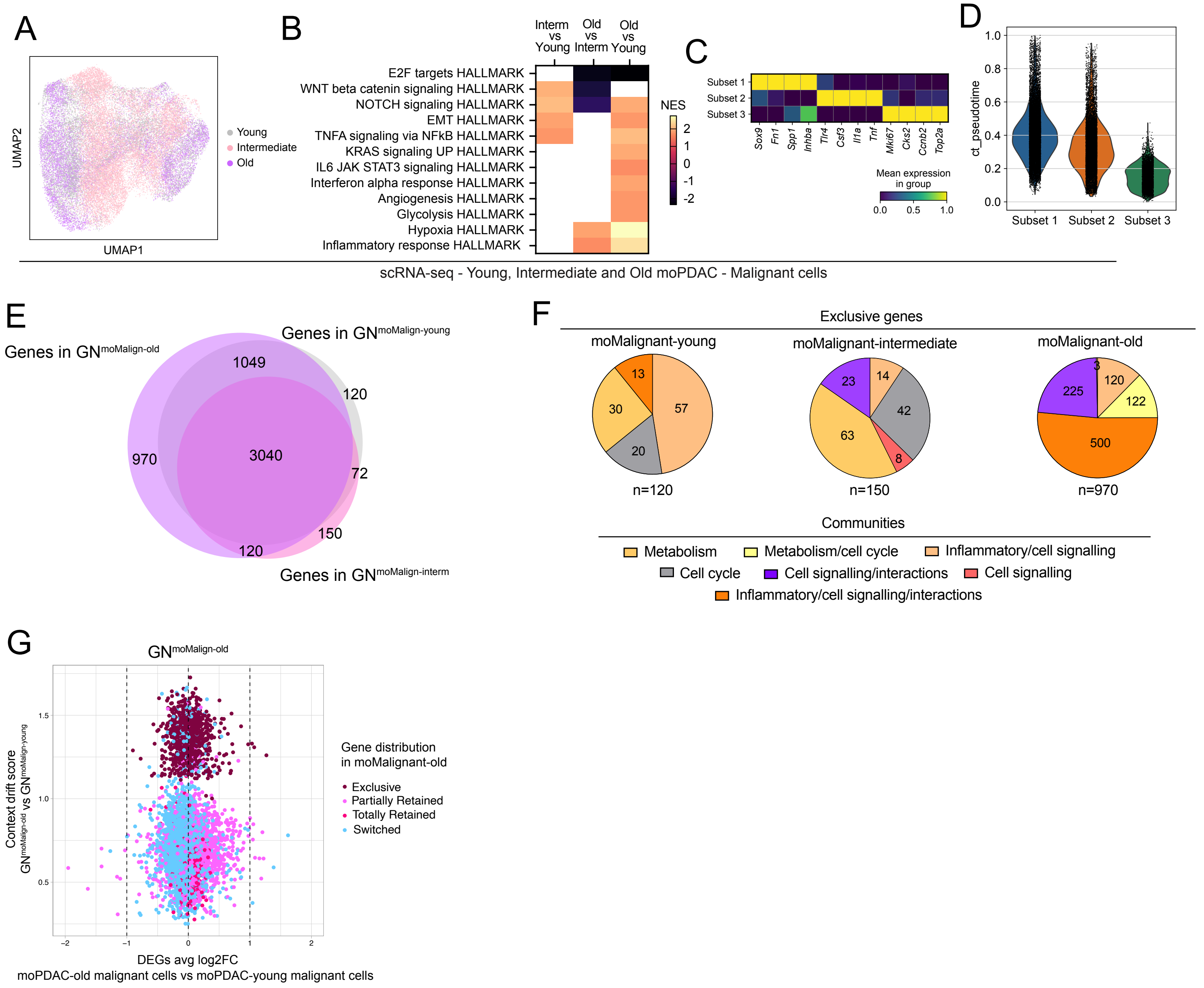


Extended Data Fig. 5

**Extended Data Figure 5. The aged TME promotes acquisition of pro-inflammatory functions in murine PDAC malignant cells. (A)** UMAP plot of malignant cells from moPDAC-young (n=8), moPDAC-intermediate (n=9) and moPDAC-old (n=8) analysed by scRNA-seq. Different ages are colour coded. **(B)** Significantly upregulated and downregulated pathways identified by GSEA of malignant cells from moPDAC-intermediate compared to moPDAC-young (left), moPDAC-old compared to moPDAC-intermediate (middle) and moPDAC-old compared to moPDAC-young (right), as assessed by MAST analysis from the scRNA-seq dataset. Blank spaces indicate when gene signatures were not significantly enriched. **(C)** Heatmap of scaled expression of selected markers in each moPDAC malignant cell sub-cluster analysed by scRNA-seq. Data are scaled such that the cluster with the lowest average expression = 0 and the highest = 1 for each gene. **(D)** Violin plots showing the distribution of ct_pseudotime scores, computed using the CellRank CytoTRACE kernel (inverse of the CytoTRACE score), across cells within each malignant subset. Subset 1 exhibits the highest ct_pseudotime values, consistent with a more differentiated state, whereas subset 3 displays the lowest ct_pseudotime values, indicative of a stem-like or less differentiated phenotype. **(E)** Venn diagram showing the overlap between the GNs of moPDAC-young malignant cells (hereon, $GN^{moMalign\text{-}young}$), moPDAC-intermediate malignant cells ($GN^{moMalign\text{-}interm}$) and $GN^{moMalign\text{-}old}$. **(F)** Pie charts showing the distribution within communities of genes 'exclusive' to $GN^{moMalign\text{-}young}$ (left, n=120), $GN^{moMalign\text{-}interm}$ (middle, n=150) and $GN^{moMalign\text{-}old}$ (right, n=970). **(G)** X-Y scatter plot of all $GN^{moMalign\text{-}old}$ genes with CDS that are DEGs (FDR < 0.05) for moPDAC-old malignant cells compared to moPDAC-young malignant cells (n=5,069), as assessed by MAST analysis of scRNA-seq data. Dotted lines mark the DEG Log2FC threshold (Log2FC > 1 or < -1). Dots (representing genes) are colour coded based on their gene distribution on $GN^{moMalign\text{-}old}$ as 'exclusive', 'switched', 'partially retained' or 'totally retained'.

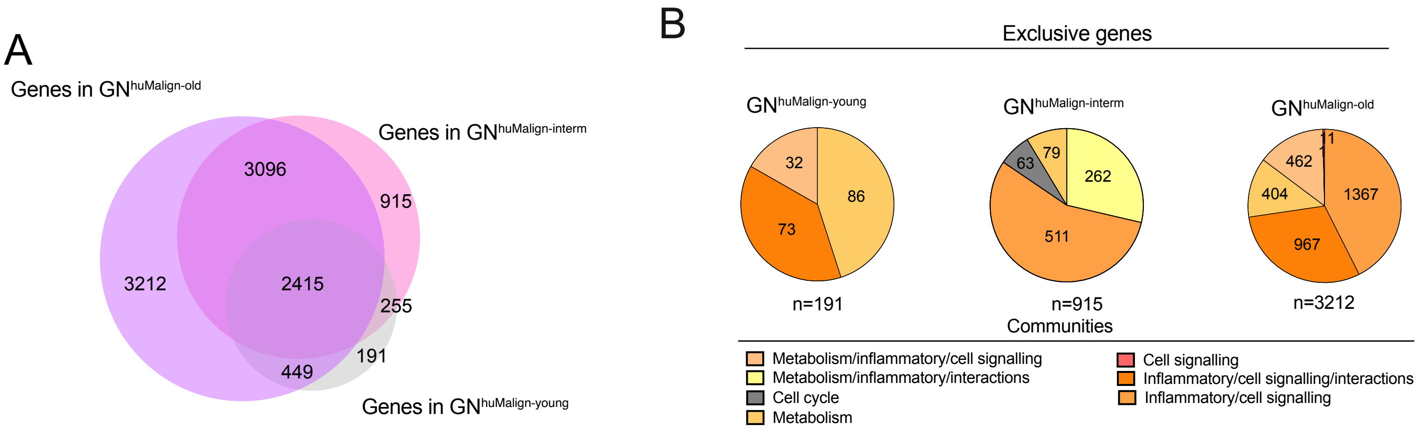


Extended Data Fig. 6

**Extended Data Figure 6. Cross-species graph-embedding highlights the aged TME as a key determinant of PDAC malignant cell biology. (A)** Venn diagram showing the overlap between genes in huPDAC-young malignant cells (hereon, $GN^{huMalign\text{-}young}$), huPDAC-intermediate malignant cells ($GN^{huMalign\text{-}interm}$) and $GN^{huMalign\text{-}old}$. **(B)** Pie charts showing the distribution within communities of genes 'exclusive' to $GN^{huMalign\text{-}young}$ (left, n=191), $GN^{huMalign\text{-}interm}$ (middle, n=915) and $GN^{huMalign\text{-}old}$ (right, n=3,212).

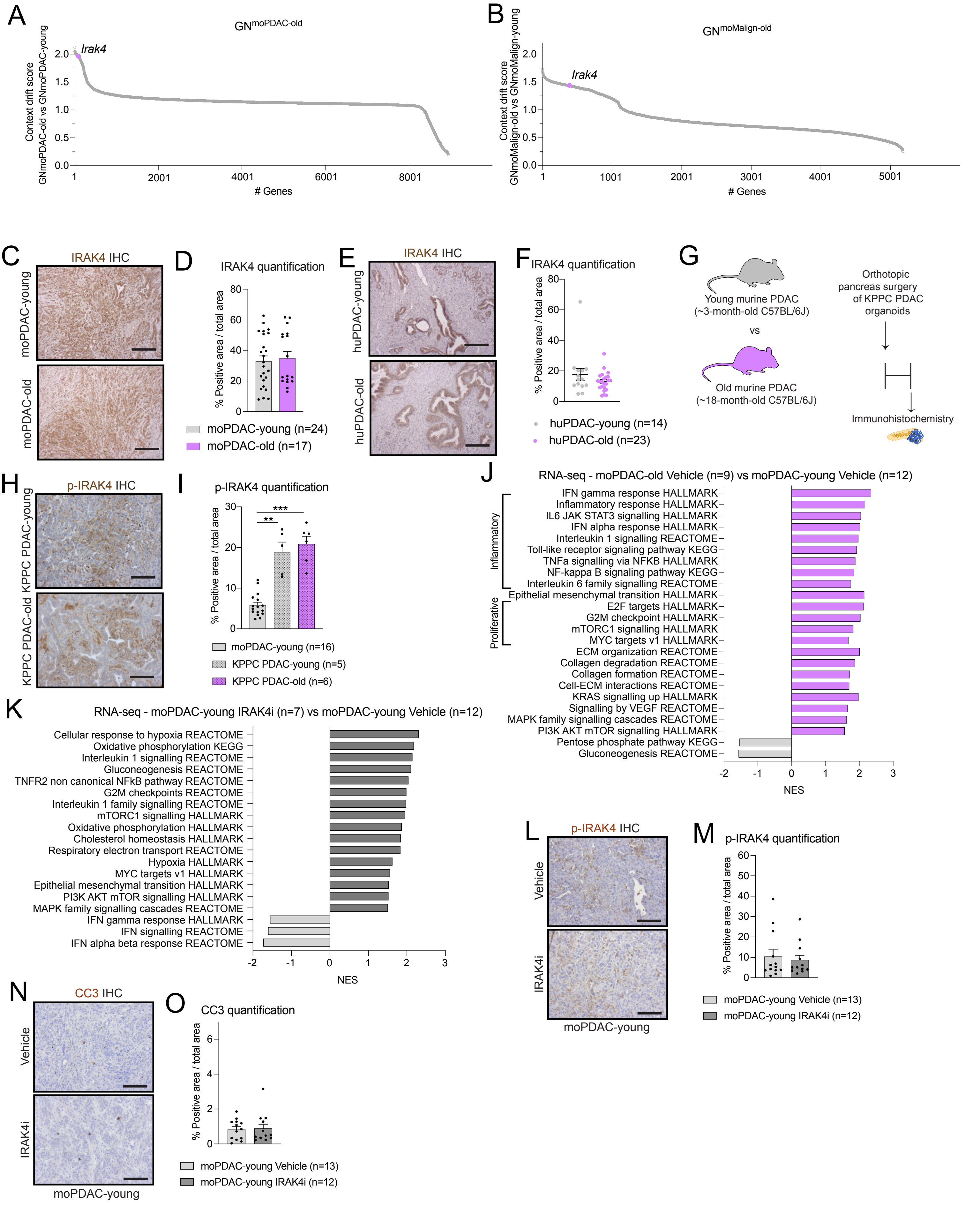


Extended Data Fig. 7

**Extended Data Figure 7. IRAK4 activity is a therapeutic vulnerability of old PDAC. (A-B)** Ranked CDS of genes on the $GN^{moPDAC\text{-}old}$ compared to $GN^{moPDAC\text{-}young}$ (with *Irak4* in position 80/8,940) **(A)** and of genes on the $GN^{moMalign\text{-}old}$ compared to $GN^{moMalign\text{-}young}$ (with *Irak4* in position 418/5,179) **(B)**. **(C)** Representative IRAK4 stains in moPDAC-young and moPDAC-old. Scale bars, 100 μm. **(D)** Quantification of IRAK4 stain in moPDAC-young and moPDAC-old. Results show mean ± SEM. No statistical difference was found, as calculated by Mann-Whitney test. **(E)** Representative IRAK4 stains in huPDAC-young and huPDAC-old. Scale bars, 100 μm. **(F)** Quantification of IRAK4 stain in huPDAC-young and huPDAC-old. No statistical difference was found, as calculated by Mann-Whitney test. **(G)** Schematic of experimental design of analyses of orthotopically-grafted murine KPPC (i.e., from $Kras^{LSL\text{-}G12D/+}$; $Trp53^{fl/fl}$; *Pdx1-Cre* GEMMs) organoid-derived PDAC models in young (~3-month-old) and old (~18-month-old) mice. **(H)** Representative phospho-IRAK4 (p-IRAK4) stains in KPPC PDAC-young and PDAC-old tumours. Scale bars, 50 μm. **(I)** Quantification of p-IRAK4 stain in (KPC) moPDAC-young, KPPC PDAC-young and KPPC PDAC-old tumours. Results show mean ± SEM. **, $P\ adj < 0.01$; ***, $P\ adj < 0.001$, Kruskal-Wallis test. **(J)** Significantly upregulated and downregulated pathways identified by GSEA of vehicle-treated moPDAC-old (n=9) compared to vehicle-treated moPDAC-young (n=12). **(K)** Significantly upregulated and downregulated pathways identified by GSEA of IRAK4i-treated moPDAC-young (n=7) compared to vehicle-treated moPDAC-young (n=12). **(L)** Representative p-IRAK4 stains in vehicle- and IRAK4 inhibitor (IRAK4i)- treated moPDAC-young. Scale bars, 50 μm. **(M)** Quantification of p-IRAK4 stain in vehicle- and IRAK4i- treated moPDAC-young. Results show mean ± SEM. No significant difference was found, as assessed by Mann-Whitney test. **(N)** Representative cleaved caspase 3 (CC3) stains in vehicle- and IRAK4i- treated moPDAC-young. Scale bars, 50 μm. **(O)** Quantification of CC3 stain in vehicle- and IRAK4i- treated moPDAC-young. No significant difference was found, as assessed by Mann-Whitney test.

### LIST OF SUPPLEMENTARY TABLES

Table S1. Metadata of human PDAC tissues used for histology analyses.

Table S2. Single cell RNA-sequencing of murine PDAC – moPDAC-old vs moPDAC-young.

Table S3. Single cell RNA-sequencing of murine PDAC – moPDAC-old vs moPDAC-intermediate.

Table S4. Single cell RNA-sequencing of murine PDAC – moPDAC-intermediate vs moPDAC-young.

Table S5. RECODR analysis of single cell RNA-sequencing of murine PDAC.

Table S6. Metadata of human PDAC tissues analysed by single cell RNA-sequencing.

Table S7. RECODR analysis of single cell RNA-sequencing of human PDAC.

Table S8. Orthologs for mouse/human PDAC all cell GN overlap.

Table S9. RECODR analysis of single cell RNA-sequencing of KPC GEMMs and orthologs for KPC/human PDAC all cell GN overlap.

Table S10. Single cell RNA-sequencing of malignant cell sub-clusters from murine PDAC.

Table S11. RECODR analysis of single cell RNA-sequencing of malignant cells from murine PDAC.

Table S12. RECODR analysis of single cell RNA-sequencing of malignant cells from human PDAC.

Table S13. Orthologs for mouse/human PDAC malignant cell-only GN overlap.

Table S14. Candidate therapeutic target prioritization strategy.

Table S15. RNA-sequencing of vehicle- and IRAK4i- treated murine PDAC.
